# Tissue-dependent mechanosensing by cells derived from human tumors

**DOI:** 10.1101/2025.01.11.632563

**Authors:** Kshitiz Parihar, Jonathan Nukpezah, Daniel V Iwamoto, Katrina Cruz, Fitzroy J Byfield, LiKang Chin, Maria E Murray, Melissa G Mendez, Anne S van Oosten, Anne Herrmann, Elisabeth E Charrier, Peter A Galie, Megan Donlick, Tongkeun Lee, Paul A Janmey, Ravi Radhakrishnan

**Affiliations:** Department of Chemical and Biomolecular Engineering, School of Engineering and Applied Science, University of Pennsylvania, Philadelphia, PA, USA; Department of Bioengineering, School of Engineering and Applied Science, University of Pennsylvania, Philadelphia, PA, USA; Institute for Medicine and Engineering, University of Pennsylvania, Philadelphia, PA, USA; Department of Physiology, Perelman School of Medicine, University of Pennsylvania, Philadelphia, PA, USA

## Abstract

Alterations of the extracellular matrix (ECM), including both mechanical (such as stiffening of the ECM) and chemical (such as variation of adhesion proteins and deposition of hyaluronic acid (HA)) changes, in malignant tissues have been shown to mediate tumor progression. To survey how cells from different tissue types respond to various changes in ECM mechanics and composition, we measured physical characteristics (adherent area, shape, cell stiffness, and cell speed) of 25 cancer and 5 non-tumorigenic cell lines on 7 different substrate conditions. Our results indicate substantial heterogeneity in how cell mechanics changes within and across tissue types in response to mechanosensitive and chemosensitive changes in ECM. The analysis also underscores the role of HA in ECM with some cell lines showing changes in cell mechanics in response to presence of HA in soft substrate that are similar to those observed on stiff substrate. This pan-cancer investigation also highlights the importance of tissue-type and cell line specificity for inferences made based on comparison between physical properties of cancer and normal cells. Lastly, using unsupervised machine learning, we identify phenotypic classes that characterize the physical plasticity, i.e. the distribution of physical feature values attainable, of a particular cell type in response to different ECM-based conditions.

## Introduction

The stiffness of normal tissues, usually characterized by a shear or Young’s modulus, varies over a wide range depending on the type of tissue, ranging from ∼100 Pa for bone marrow or fat to near MPa for some muscles and even more for cartilage^1,2^. For a specific tissue, animal, and age, the elastic modulus is a tightly controlled quantity, but it undergoes large changes in some diseases such as fibrosis and cancer, especially when the tissues are mechanically stressed as they are *in vivo*^3^. The effects of changes in the mechanical properties of tissues on the cells within them are increasingly implicated both in the normal development of tissues and in the pathological functions of cells in various disease states^4–6^. In addition to changes in physical properties, changes in tissue stiffness triggered by injury or disease can lead to chemical changes in the extracellular matrix (ECM) including its most abundant constituents such as collagen, fibronectin, and hyaluronic acid. In cancer biology, the increased stiffness of several types of tumors, most prominently breast, liver, and colorectal tumors, has motivated studies of mechanical sensing by both cancer and stromal cells, which have yielded accruing evidence that changes in cellular environment both in vitro and in vivo can affect the structure and function of cells in a manner that is promotes malignancy^7,8^. The combination of physical and chemical changes in ECM can direct tumor growth and dispersion^9^. Apart from altered ECM conditions in the primary tissue site, metastasizing cancer cells will often encounter substantially different ECM chemical composition and mechanical stiffness in metastatic tissue sites (such as lung to brain, breast to bone metastasis^10^). Therefore, it is crucial to understand the response of the cells to changes in ECM stiffness and chemistry.

Precisely how cells detect or respond to mechanical signals, and how that response depends on the chemistry of the ECM is only beginning to be revealed, with many proteins implicated in the capacity of cells to respond to mechanical cues^11^. It is also not clear whether a response to mechanical cues is a general feature of most cell types or whether it is different in normal and malignant cells. Studies have found differing responses to mechanical environments even among closely related cell types^12^. For example, pluripotent cells derived from human bone marrow can be directed to differentiate into highly distinct cell types by manipulating their substrate elastic modulus to levels that span the range from very soft tissues like the brain to much stiffer tissues like muscle or bone^13^. In contrast, a similar study of murine embryonic stem cells showed very little dependence of the cells on substrate stiffness^14^. In the context of cancer, a reasonable expectation might be that since one of the hallmarks of the malignant transformation is the ability to grow in soft agar^15^, cancer cells might have much less response to mechanical signals than the normal cells from which they derived, which typically are much more limited in their proliferation rates on substrates with normal physiological stiffness^16^. One of the earliest quantitative studies of stiffness sensing in fibroblasts showed that whereas NIH 3T3 fibroblasts responded very strongly to changes in substrate elastic modulus, by altering their morphology, their cytoskeletal assembly, and the degree of protein tyrosine phosphorylation, transfection of these cells with H-Ras eliminated the response of the cells to stiffness, producing a phenotype similar to that seen on plastic even when cells were grown on very soft substrates^17,18^. However, a total loss of mechanical sensitivity is not a universal feature of cancer cells^11,12^, and even highly abnormal cells such as HeLa cells respond robustly to changes in substrate elastic modulus^19,20^.

Thus, to gain a perspective on the relative importance of substrate characteristics on the structure, and motility of cancer cells on substrates with defined stiffness and chemical composition, 25 different human cancer cell lines derived from tumors in breast, colon, brain, ovary, pancreas, prostate, and skin tissues were studied along with 5 immortalized but non-tumorigenic cell lines from the same set of tissues. Using this pan-cancer mechanobiology dataset, we inferred how the cellular mechanics (adherent area, cell shape, cell stiffness, and motility) are influenced by varying biologically realistic ECM physicochemical properties that capture cell-ECM interactions for major solid tumor types. We find heterogeneity to be the norm in how different cell types respond in terms of change in cellular mechanics to mechanosensitive and chemosensitive changes in ECM conditions. Our results also caution against generalizing comparison of normal vs cancer cell mechanics across and even within tissue types. Lastly, we show that behavior of different cell lines across various substrates can be grouped into physical feature-specific classes that define cellular plasticity in terms of the range of physical feature values a specific cell line can attain in response to a specific substrate.

## Results

The pan-cancer Leidos dataset provides a rich array of cellular physical measurements (spread area, shape (circularity and aspect ratio), cell stiffness, and motility) across 30 cell lines spanning 8 different tissue types on varying substrates which consist of polyacrylamide (PAAm) gels with elastic moduli of 500 Pa or 30 kPa coated with either fibronectin (500 Pa FN, 30 kPa FN) or collagen I (500 Pa Coll, 30 kPa Coll), cross-linked hyaluronic acid gels with elastic modulus of 500 Pa or less coated with fibronectin (HA FN) or collagen I (HA Coll), or glass slides (Fig. 1A). Out of the 30 cell lines, 25 are cancer cell lines and 5 are immortalized but non-malignant (normal) cell lines (Fig. 1D). Live cell imaging at 24 hours was used to measure cell area, aspect ratio, and circularity. Cell stiffness was assessed using atomic force microscopy, yielding measurements of Young’s modulus. Motility was quantified by tracking individual single cells during time lapse movies. For example, figure 1B shows the snapshot images from brightfield microscopy for T98G cell line along with median values of its measured physical features across different substrates. Analyzing the relationship between the physical features, we found cell area to be highly correlated with shape features (aspect ratio and circularity) (Supp. Fig. 1). Cell stiffness is not correlated with any of the other physical properties, and motility is only weakly related to both morphology and cell stiffness (Supp. Fig. 1).

**Figure 1:**
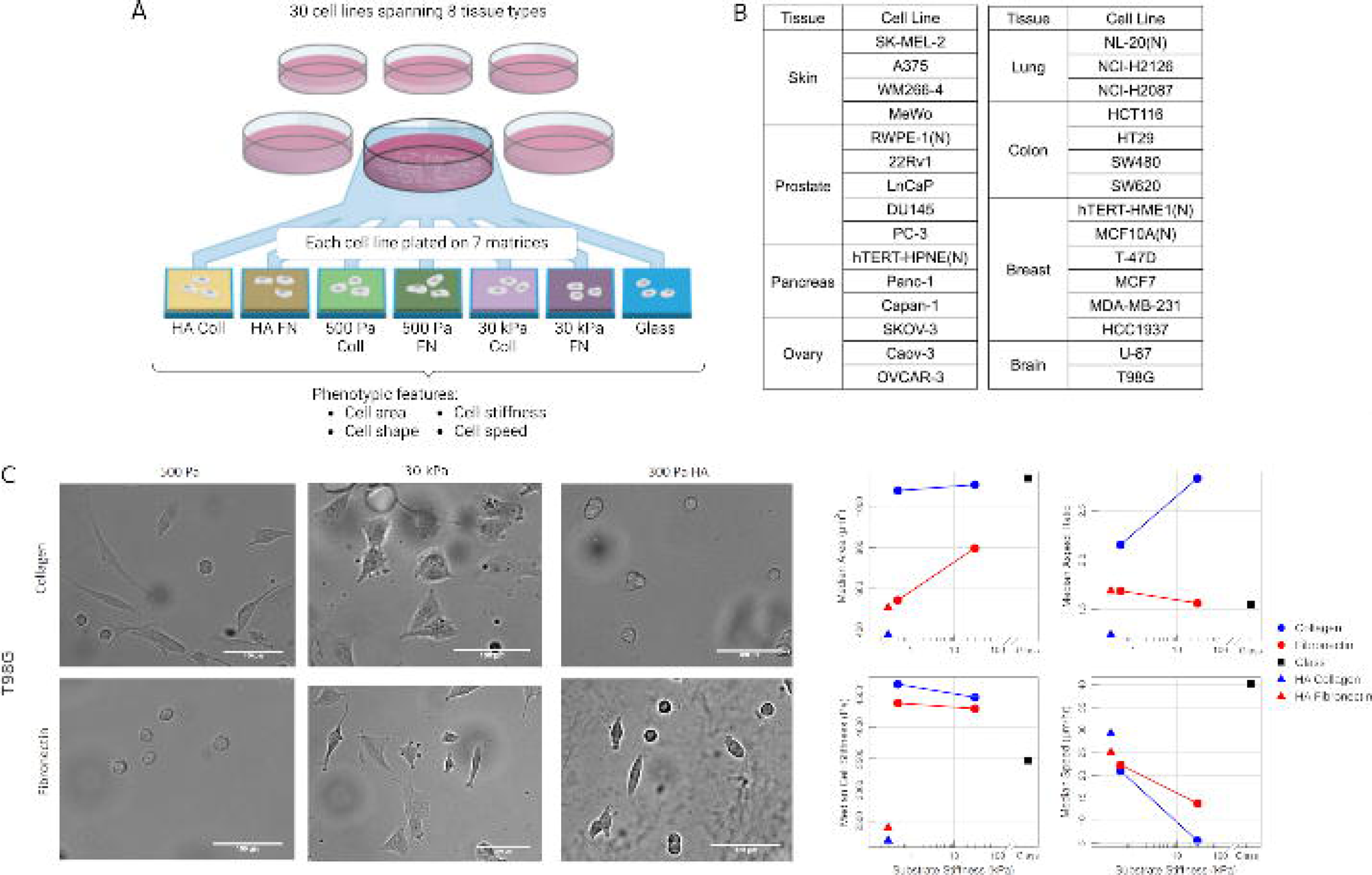
**(A)** 30 cell lines (including 25 cancer and 5 normal cell lines) that span 8 different tissue types were grown on 7 different substrates and physical properties of the cells (namely, area, shape (circularity and aspect ratio), stiffness, and motility) were measured^92^. 500 Pa Coll, 500 Pa FN, 30 kPa Coll, 30 kPa FN: polyacrylamide (PAAm) gels with elastic moduli of 500 Pa or 30 kPa coated with either fibronectin or collagen I. HA Coll, HA FN: cross-linked hyaluronic acid gels with elastic modulus of 500 Pa or less coated with fibronectin or collagen I. **(B)** Tables listing the tissue type and cell lines considered. (N) refers to non-malignant (normal) cell lines. Examples of cell area, aspect ratio, stiffness, and motility (median values) on different substrates for T98G along with phase-contrast image of the cells.

### I. Heterogeneity in sensitivity to changes in ECM stiffness and composition

One of the primary ways cells respond to ECM alterations is through changes in their physical properties. We thus asked, how sensitive are the morphology, cell stiffness, and motility of different cell lines to change in substrate conditions? The effect of substrate sensitivity on the phenotypic features is analyzed using the relative change in the median value of the feature when moving from one substrate to another. Due to skewness observed in the data, we used the median value as a population-level representative of a cell line’s physical behavior on a particular substrate. We analyze both mechanosensitive and chemosensitive changes in ECM conditions. The alterations considered include (i) increase in stiffness of PAAm-based substrate (30k-500Pa Coll and 30k-500Pa FN), (ii) 500 Pa PAAm-based to 500 Pa HA-based substrates (HA-500Pa Coll and HA-500Pa FN), (iii) change in integrin ligand on HA-based substrate (HA Coll-FN), on soft (500Pa Coll-FN) and stiff (30kPa Coll-FN) PAAm-based substrates, (iv) 500 Pa HA-based to stiff 30 kPa PAAm-based substrate (30kPa-HA Coll and 30kPa-HA FN) and (v) stiff PAAm-based substrate to extremely high, supraphysiological stiffness of glass (Glass-30kPa Coll and Glass-30kPa FN).

Increase in ECM stiffness of PAAm-based substrate (30k-500Pa) significantly increases area in melanoma cells, and a similar effect is also observed with a soft (500 Pa) HA substrate (HA-500Pa) (Fig. 2A). Combined with results from 30kPa-HA Coll and 30kPa-HA FN (Supp. Fig. 1A), we can see that melanoma cells can spread more on a soft HA ECM as compared to a stiff ECM. In contrast, change from 500 Pa PAAm to 500 Pa HA based substrate (HA-500Pa) causes a decrease in area for several cell lines in other tissue types (Fig. 2A). Interestingly, even on stiff PAAm-based substrate, some of the cell lines (22Rv1, SW620, U-87, Capan-1, and SKOV-3) show a decrease in spread area as compared to soft PAAm-based substrate (30k-500Pa, Fig. 2A). Corresponding to an increase or decrease in area with change in ECM conditions, the shape of the cell becomes more elongated (higher AR or lower Circularity) or more rounded (lower AR or higher Circularity) respectively (Fig. 2B, Supp. Fig. 2B, 3).

**Figure 2:**
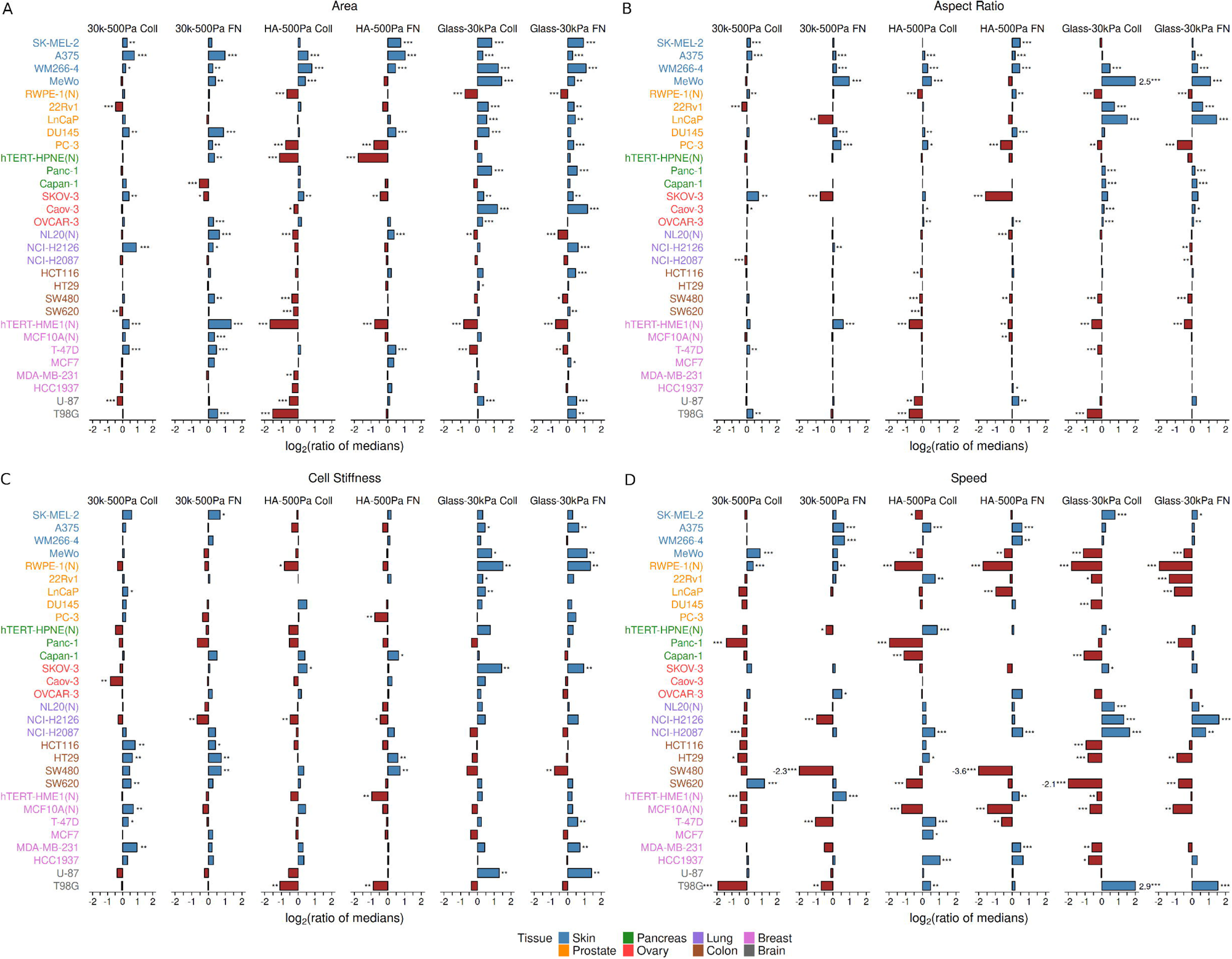
For each cell line, the ratio of the median values for **(A)** area, **(B)** shape (aspect ratio), **(C)** cell stiffness, and **(D)** cell speed as a measure of the phenotypic sensitivity to substrate change. (30k-500Pa Coll: 30 kPa Coll/500 Pa Coll, 30k-500Pa FN: 30 kPa FN/500 Pa FN, HA-500Pa Coll: HA Coll/500 Pa Coll, HA-500Pa FN: HA FN/500 Pa FN, Glass-30kPa Coll: Glass/30 kPa Coll, Glass-30kPa FN: Glass/30 kPa FN). (N) refers to non-malignant (normal) cell lines. See also supplementary figures 2 and 3. ***p-value < 0.01; **p-value < 0.05; *p-value < 0.1, adjusted for multiple testing using Benjamini-Hochberg procedure. For each cell line on a particular substrate, *n* > 25 cells. See supplementary tables 1-3 for the exact value of *n* for the cell lines.

In contrast to the substrate effects on cell area, the stiffness of melanoma cells does not change significantly with changes in ECM stiffness or in presence of HA (Fig. 2C). Colon cells, on the other hand, show an increase in cell stiffness on stiffer PAAm-based substrates (30k-500Pa) as well as for HA-500Pa FN alteration (Fig. 2C). Notably, some of the cell lines also show a decrease in cell stiffness with increasing ECM stiffness (Fig. 2C). As compared to changes observed in area, a switch in the type of integrin ligand (HA Coll-FN, 500Pa Coll-FN, 30kPa Coll-FN) does not have significant effect on cell stiffness (Supp. Fig. 2A, C). Going from gel-based substrates to glass leads to a significant increase in cell stiffness across several cell lines (Fig. 2C), however even in this case the opposite behavior of decreasing cell stiffness can be seen.

In contrast to cell stiffness, more cell lines undergo significant changes in cell motility with change in substrate composition from 500 Pa PAAm-based to 500 Pa HA-based substrate (HA-500Pa) (Fig. 2D) and change in ECM ligands (HA Coll-FN, 500Pa Coll-FN, and 30kPa Coll-FN) (Supp. Fig. 2D). Increase in ECM stiffness (30k-500Pa) can lead to either increase or decrease in migration speeds in a cell line dependent manner across tissue types (Fig. 2D). A similar mix of increase and decrease in motility can also be observed in the switch from gel-based substrate to glass.

Altogether, there is substantial heterogeneity in the mechanosensitive and chemosensitive response of cell lines across and within tissue types to changes in ECM composition and stiffness. In particular, the changes in different physical properties (morphology, cell stiffness, and motility) vary considerably compared to each other.

### II. Difference between normal and cancer in terms of physical response depends on the substrate conditions and the cell lines being compared

Comparing the cell lines in terms of the median and distribution of the physical feature values across all the 7 different substrates shows some intriguing differences between cancer and normal cell lines. The normal breast cell lines are stiffer as compared to their cancer counterparts, which is in contrast to the normal lung cell line being softer than cancer cells (Fig. 3B). The normal cell lines (namely, hTERT-HPNE, NL20, and hTERT-HME1) have greater median migration speeds and a much broader distribution than their cancer counterparts in the respective tissue types (Fig. 3C). Other observations to note are that the pancreatic normal cell line hTERT-HPNE has a much larger area and highly non-circular shape (i.e. high aspect ratio) (Fig. 3A, Supp. Fig. 4A), and the pancreatic cancer cell line Capan-1 has much greater cell stiffness values as compared to all the other cell lines (Fig. 3B, Supp. Fig. 4B).

**Figure 3:**
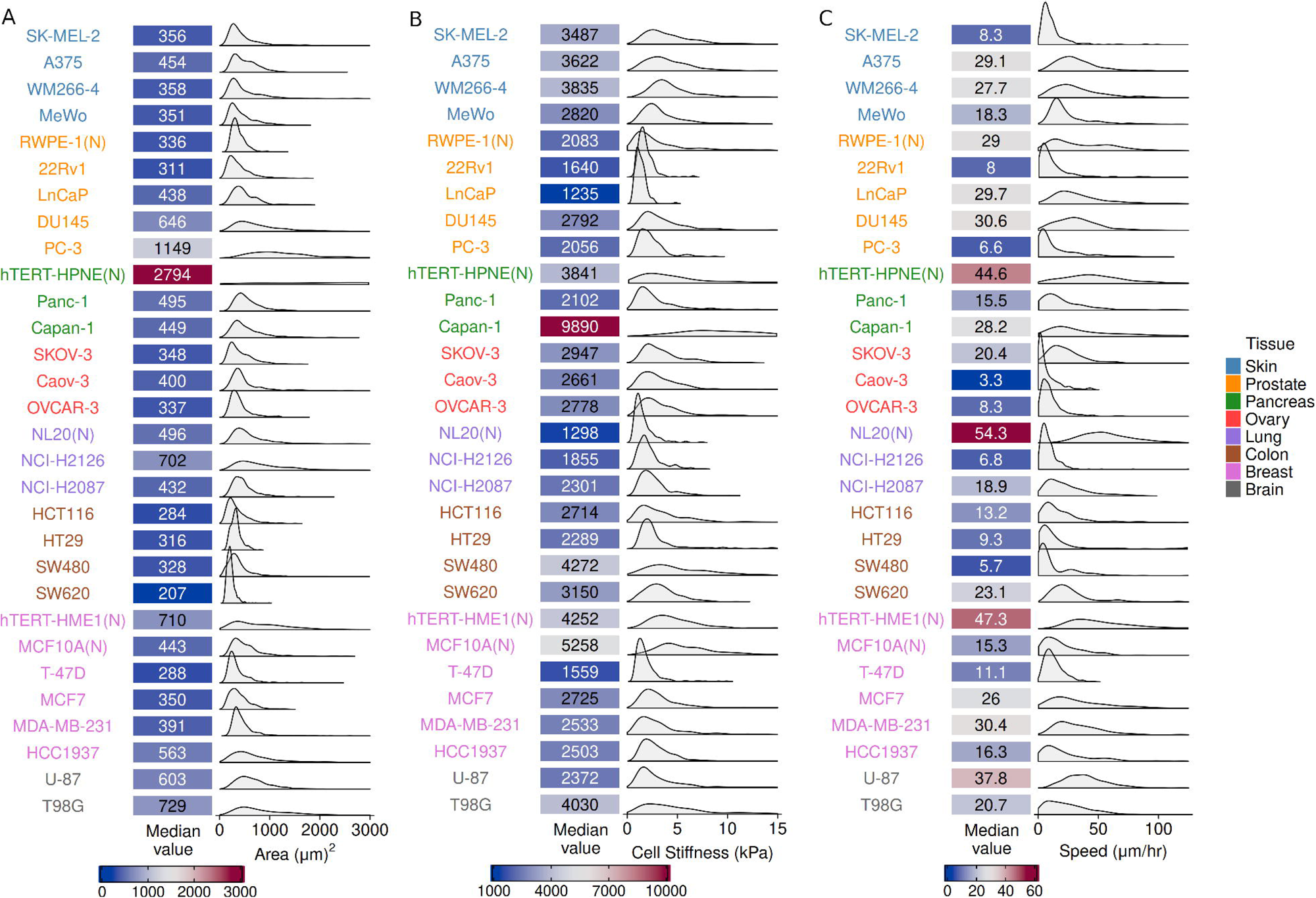
Median values of cell **(A)** area, **(B)** stiffness, and **(C)** speed across all the 7 different substrates, along with the kernel density estimates (KDE) for summarizing the distribution of feature values for each cell line. (N) refers to non-malignant (normal) cell lines. See also supplementary figure 4. Measurements from *n* > 200 cells for each of the cell lines. See supplementary tables 1-3 for the exact value of *n* for the cell lines.

Next, we assessed how the cancer and normal cell behavior differ within each of the substrate types by comparing the median values of the physical features (Fig. 4). The normal breast cell line hTERT-HME1 has greater median area than all the cancer cell lines on 500 Pa Coll and 30 kPa PAAm substrates but not on HA substrates (Fig. 4D). The other normal breast cell line MCF10A, on the other hand, has median area smaller than or similar as HCC1937 on all substrates and greater median area than the other 3 cancer cell lines on 500 Pa FN, 30 kPa, and glass substrates (Fig. 4D). Similar cell line dependence in cancer vs normal comparison can be seen for prostate and lung tissues as well (Fig. 4A,C). The pancreatic normal cell line hTERT-HPNE has substantially higher spread area values, not only compared to other pancreatic cell lines but all the other cell lines, on 500 Pa PAAm, 30 kPa PAAm, and glass substrates with reduced though still high area on HA substrates (Fig. 4B). In the context of aspect ratio, pancreatic, lung, and breast normal cell lines have higher aspect ratio than the corresponding cancer cell lines on 500 Pa and 30 kPa PAAm substrates (with a few exceptions) (Supp. Fig. 5B-D). In contrast, the prostate normal vs cancer cell line comparison of aspect ratio is much more cell line dependent (Supp. Fig. 5A).

**Figure 4:**
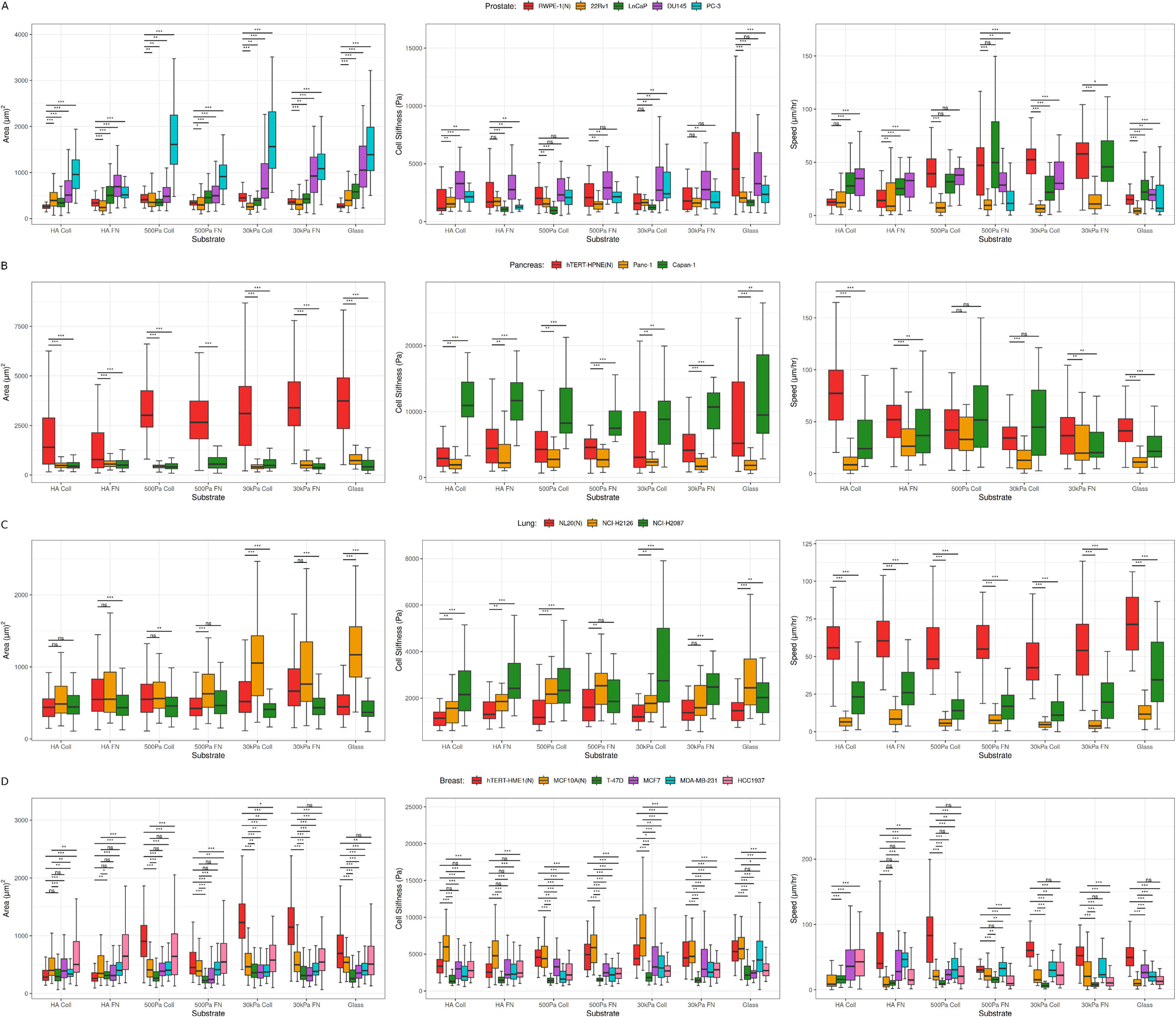
Comparing tissue-specific normal and cancer cell behavior in terms of spread area, cell stiffness, and motility for **(A)** prostate, **(B)** pancreas, **(C)** lung, and **(D)** breast cell lines on soft (500 Pa) and stiff (30 kPa) PAAm Coll and FN substrates, soft (500 Pa) HA substrates coated with Coll and FN, and glass. (N) refers to non-malignant (normal) cell lines. See also supplementary figure 5. ***p-value < 0.01; **p-value < 0.05; *p-value < 0.1, adjusted for multiple testing using Benjamini-Hochberg procedure. For each cell line on a particular substrate, *n* > 25 cells. See supplementary tables 1-3 for the exact value of *n* for the cell lines.

In terms of cell stiffness, all the breast cancer cell lines are softer than MCF10A on all the substrates and softer than hTERT-HME1 on all except HA substrates (Fig. 4D). In contrast, the lung cancer cell lines are stiffer than the normal cell line NL20 on all the substrates (Fig. 4C). For prostate tissue, similar to area and AR, the cancer vs normal comparison with cell stiffness as the metric is cell line dependent on all substrates with the exception of HA Coll where RWPE-1 has lower median cell stiffness than the cancer cell lines (Fig. 4A). In the case of pancreas, the outlier-like behavior of Capan-1 cancer cell line in terms of cell stiffness as compared to other cell lines (pancreatic and other tissues) can be seen on all the substrates (Fig. 4B).

For cell motility as the metric of response, the normal lung cell lines NL20 is substantially faster than the cancer cell lines on all the substrates (Fig. 4C). While the normal prostate cell line RWPE-1 is faster than non-metastatic cancer cell line 22Rv1 across the substrates, RWPE-1 has higher median speed than metastatic cell lines^21,22^ (LNCaP and DU145) only on 30 kPa matrices (Fig. 4A). Pancreatic normal cell line hTERT-HPNE shows higher median migration speed than both the cancer cell lines only on the HA Coll, 30 kPa FN, and glass substrates (Fig. 4B). For breast tissue, hTERT-HME1 normal cells are faster than all the cancer cells on 500 Pa Coll, 30 kPa and glass substrates, whereas comparison of cancer cell lines with MCF10A are much more cell line dependent on all the substrates (Fig. 4D).

Altogether, the results show that for each physical feature as the metric of response there is a lack of generalization between tissue types for comparisons between normal and cancer cells on all the substrates considered. Within a tissue type, there are cases of both cell line independent (like normal lung cancer cells being faster than cancer cells, breast normal cells being softer than cancer cells) and cell line dependent comparisons between normal and cancer cells for metric of response being any of the physical features.

### III. Distinct mechanotypes characterize the physical response of cells to ECM-based cues

The highly varied response of cell lines observed in terms of sensitivity to changes in substrate conditions led us to ask if there are potential underlying similarities in how different cell lines phenotypically behave on different substrates. We first performed principal component analysis (PCA) using the median values of the five physical features for each of the cell line-substrate pairs (Sup. Fig. 6). There were no clear clusters in the PCA plots except the separation of outlier-like cases mentioned in Section II for area and cell stiffness, namely hTERT-HPNE (normal, pancreas) and Capan-1 (cancer, pancreas) respectively. This suggests that the potential underlying similarities in phenotypic behavior across substrates might be physical feature-specific instead of being defined by a combination of them.

Next, we used all the measurements (instead of only median values as measure of response) to compare physical feature-specific behavior of cell lines across the substrates. Figure 5A shows the analysis workflow employing unsupervised machine learning framework using consensus clustering^23,24^ to identify phenotypic classes for each physical behavior. For comparison of cell lines within and across substrates (i.e. comparing cell line-substrate pairs), we used Wasserstein-1 distance^25,26^. The number of phenotypic classes for each of the physical features is determined based on the proportion of ambiguous clustering (PAC)^27^ and Calinski-Harabasz Index (CHI)^28^. That is, the optimal number of clusters would correspond to low PAC and high CHI. Additionally, given the inherent uncertainty in unsupervised clustering, we also separate out possible boundary cases (cell line-substrate pairs) which do not have a high confidence score for being part of one cluster over another (see methods for details).

**Figure 5:**
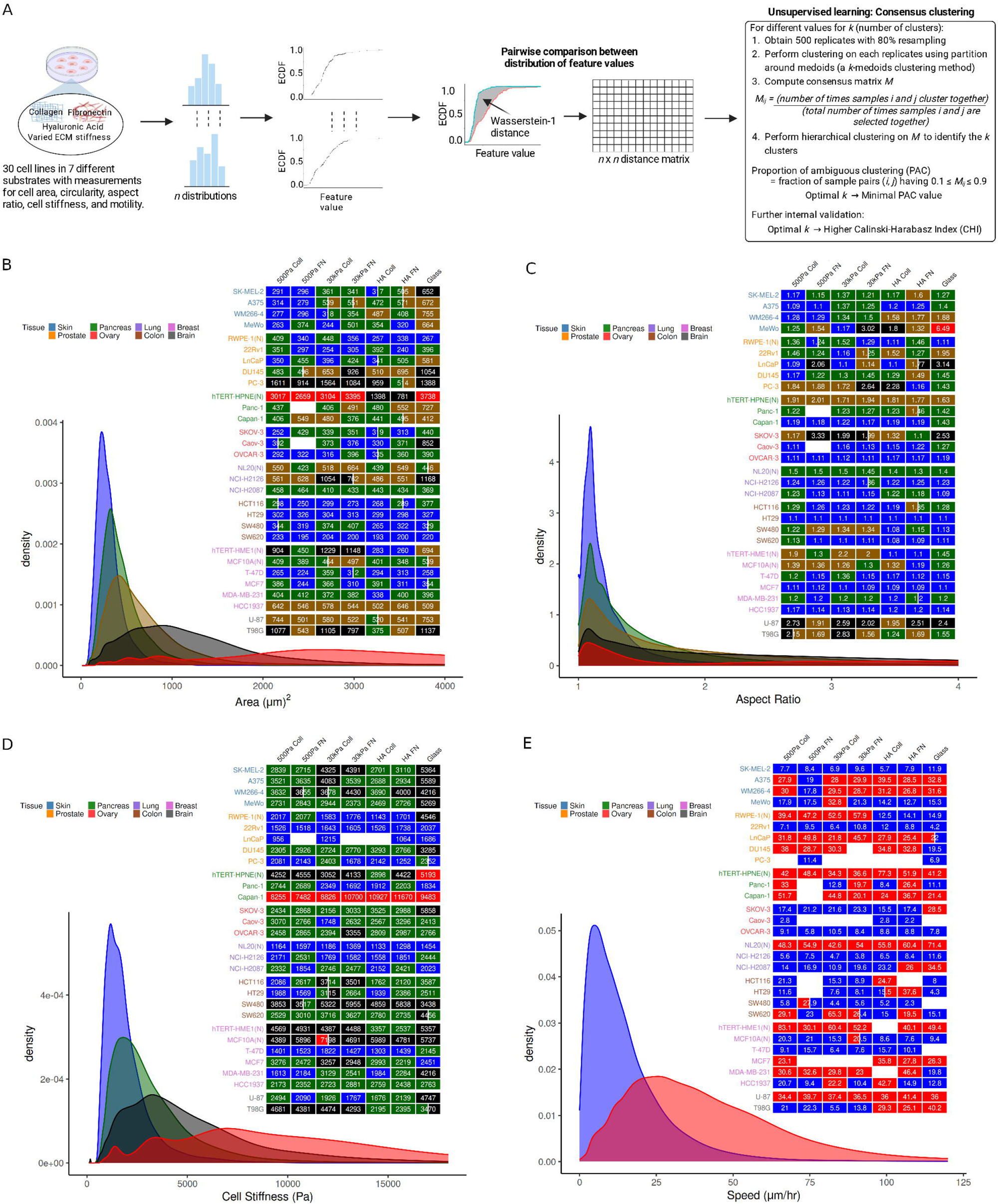
**(A)** Schematic representation of unsupervised learning approach to identify phenotypic classes (mechanotypes) within each of the physical behaviors^92^. (ECDF: Empirical cumulative density function). Mechanotypes for cell **(B)** area, **(C)** aspect ratio, **(D)** stiffness, and **(E)** motility, whereby the heatmaps show the phenotypic class for each cell line-substrate pair and the KDEs correspond to characteristic density function for each class. The numeric values shown in the heatmaps correspond to median values of the physical feature for each cell line-substrate pair. (N) refers to non-malignant (normal) cell lines. See also supplementary figures 7 and 8. Note that in this analysis only those cell line-substrate pairs which have at least 25 data points for the physical feature of interest are considered.

Based on this unsupervised learning methodology, 5 classes (mechanotypes) for area and aspect ratio, 4 for cell stiffness, and 3 and 2 mechanotypes respectively for circularity and motility were identified (Fig. 5B-E, Supp. Fig. 7,8). The outlier-like cases of area (hTERT-HPNE) and cell stiffness (Capan-1) noted in Section II separate out into a distinct class. Also, MeWo on glass is separated out as a single class for aspect ratio due to its much higher values as compared to other cell line-substrate pairs. Thus, we primarily have 4 mechanotypes each for area and aspect ratio, and 3 mechanotypes for cell stiffness. We also compared the performance of Kolmogorov-Smirnov (KS) distance, a commonly used metric for comparing distributions, in identifying relevant clusters. As compared to Wasserstein-1 distance, unsupervised clustering based on KS distance was unable to separate out the outlier-like behavior of hTERT-HPNE in area and Capan-1 in cell stiffness (Supp. Fig. 9).

The difference between mechanotypes can be seen from their respective kernel density estimates (KDEs), obtained by combining data from all the cell line-substrate cases falling in that class (the boundary cases have not been included in the density estimation). More specifically, we can see that certain cell lines have more plasticity in terms of the range of values they can attain for a specific physical feature as compared to others on a particular substrate and across substrates. Each substrate has at least one cell line within each of the mechanotypic classes for each physical feature. Within each tissue type, there is a mix of 2 or more mechanotypes with some interesting patterns. Among breast tissue cell lines, the normal cell lines (hTERT-HME1 and MCF10A) and highly aggressive basal-like cell lines^29,30^ (MDA-MB-231 and HCC1937) mainly have intermediate to high area mechanotypes, whereas less aggressive luminal-like cell lines^29,30^ (T-47D and MCF7) primarily have the lowest area mechanotype (Fig. 5B). The colon-derived cells are largely part of only the lowest area mechanotype (blue KDE in Fig. 5B) and belong to the intermediate to high cell stiffness mechanotypes (green and black KDEs in Fig. 5D). Similarly, ovary cells primarily show an intermediate cell stiffness mechanotype (green KDE in Fig. 5D) along with lower area mechanotypes (blue and green KDEs in Fig. 5B). Skin cells, on the other hand, show intermediate to high cell stiffness mechanotypes, but with a more varied range of area mechanotypes as compared to colon and ovary cells. In contrast to these tissues, the low cell stiffness mechanotype (blue KDE in Fig. 5D) is primarily present in lung and prostate tissues.

It is important to note the possibility that a cell line may stay within the same mechanotypic class across two substrate conditions but change its primary sampling region within the distribution leading to a significant change in physical property. For instance, SW620 stays within the low area (blue KDE in Fig. 5B) and intermediate cell stiffness (green KDE in Fig. 5D) mechanotype in both 500 Pa Coll and 30 kPa Coll but shows a significant decrease in median area and increase in median cell stiffness with an increase in ECM stiffness from 500 Pa Coll to 30 kPa Coll (Fig. 1A, C).

In summary, the unsupervised learning-based analysis shows that the versatility in the physical response of cells to ECM composition and stiffness can be categorized into specific classes (mechanotypes) mapping to narrow, intermediate, and broader distributions of physical feature values. To alter specific aspects of cellular mechanics with change in substrate conditions, the cells can stay within the mechanotype but change the region they predominantly sample from in the corresponding distribution, or simply switch between mechanotypes.

## Discussion

Cellular function, normal or pathological, is influenced by interactions with the microenvironment^4^. Here, we studied how differing ECM mechanics and chemical compositions impact the physical response of cancer and normal cell lines from 8 different tissue types. Conventionally, it has been expected that cells will spread more and increase in stiffness with increasing substrate stiffness. However, accruing evidence suggests a more complicated picture^12^. While increase in spread area on stiffer ECM has been previously shown for fibroblasts^31^, endothelial cells^32–35^, hepatic stellate cells^36–39^, and myocytes^40–44^, insensitivity to change in ECM stiffness or a decrease in area on stiffer ECM has also been reported for several other cell types^14,45–48^. These cellular responses are further contingent on the type and density of the integrin ligand available for adhesion^49–51^. In the context of cell stiffness, similar contrasting behaviors of increase in cell stiffness or insensitivity to change in ECM stiffness depending on the cell type have also been reported^13,14,50,52–54^. In the case of motility, it has been argued that the migration speed of cells has a maximum speed at a particular substrate stiffness and thus, speed can increase or decrease with changing ECM stiffness depending on which side of the peak speed cell is on^55–57^. The ECM stiffness for peak speed appears to differ among cell types and is further dependent on the ligand density^55,57^. Specifically for cancer cells, studies have found differing physical responses to change in substrate stiffness^17,45,54,58–60^. Consistent with these prior studies, our results of 30 distinct cancer and normal cell lines also show a diverse trend within and across tissue types in mechanosensitive response to increasing ECM stiffness, with the changes in morphology, cell stiffness, and motility further dependent on integrin ligand type. While the conventional picture of increasing area and cell stiffness could be seen more consistently across cell lines in the case of going from gel-based matrix to extremely high, supraphysiological stiffness of glass, even in this case there were some exceptions.

Hyaluronic acid (or hyaluronan, HA), a soft polymeric glycosaminoglycan present in ECM, plays an important role in development, wound healing, and cancer progression^61–64^. Prior studies have found that myocytes, fibroblasts, glioma, and hepatocellular carcinoma cells can respond to soft HA matrices coated with integrin ligands similar to stiff substrates in terms of increase in spread area and cell stiffening^50,65–67^. Importantly, these studies also found that the response was integrin ligand dependent and not universal across cell types^50,66,67^. To ascertain whether this extends to other cell types, we analyzed how the physical response of cancer and normal cell lines change on integrin ligand (fibronectin or collagen I) coated soft HA-based substrates as compared to soft and stiff inert PAAm-based substrates. Most melanoma cell lines responded with as much or even more increase in spread area on soft HA-based substrates as compared to stiff PAAm-based substrates. However, similar to mechanosensitivity, overall there was heterogeneity in the response (insensitive or significant increase/decrease) to the presence of HA across cell lines and integrin ligand types for all the physical features (area, shape, cell stiffness, and motility). While the observed effect of HA to alter cell response is not universal, our results nevertheless emphasize an important aspect, often not reported in previous studies, of considering the role ECM mechanics in conjunction with HA when studying how a particular cell type may respond in vivo.

Mechanistic understanding of how the presence of HA, especially with an integrin ligand, impacts cellular mechanics is sparse. HA interacts with cells through transmembrane receptors, like CD44 and RHAMM, and these linkages can regulate cell adhesion and motility^66,68,69^. Prior studies suggest potential augmentation of integrin-mediated signaling with HA-mediated signaling leading to cellular response on soft matrix otherwise only seen on stiff matrix^50,66,67^. In particular, Mandal et al. showed that hepatocellular carcinoma cell line (Huh7) spread as much on soft HA matrices as on stiff matrices and that decreasing PIP_2_ levels or inhibiting PIP_2_ or PIP_3_ function reduced spreading and adhesion of cells to HA but not on stiff substrates^67^. Combined with the reports that proteins involved in actin polymerization, such as formins^70,71^ and N-WASP^72^, and focal adhesion proteins, such as alpha-actinin^73^ and vinculin^73,74^, can be activated by either force or binding phosphoinositides, cells adhering to soft substrates containing both an integrin ligand and HA may form focal adhesions and undergo cytoskeleton reorganization (similar to that observed on stiff substrates) depending on the synthesis and spatial distribution of phosphoinositides. More work is thus needed to understand the mechanisms underlying the interplay between HA and integrins in mediating cellular mechanics across different cell types.

Tissue specific comparisons of cancer and normal cell lines showed that for each of the physical features as a metric of response the differences are cell line specific rather than general. Some interesting trends though can be observed at the tissue level. Among breast cell lines, normal cells were consistently stiffer than cancer cells on all substrates except on HA matrices where one normal cell line (MCF10A) remained stiffer but other normal cell line (hTERT-HME1) had comparable cell stiffness with cancer cells. This is consistent with a prior study which found cancer cells derived from breast, bladder, cervix, and pancreatic tumors to be softer than their normal counterparts^45^. However, it is important to note that the study used collagen-coated glass coverslip as opposed to the much broader range of substrate types used in our analysis^45^. Further, in contrast to breast, normal lung cells were softer than their cancer counterparts across different substrate types. These normal lung cells also had much higher migration speeds than cancer cells on all the substrates. In breast, however, one normal cell line (hTERT-HME1) had higher motility than cancer cell lines on several but not all substrates, whereas the other normal cell line (MCF10A) had similar or lower speeds than cancer cells. Altogether, the results show that to study differences/similarities between cancer and normal cells using physical properties as the metric, it is important to take into account both substrate conditions (mechanical and chemical) and the cell types being compared.

Most prior studies rely on gauging differences in cellular behavior on different substrates by comparing the sample mean or median values of the metric of response. Their behavior can also be analyzed more holistically, by comparing the distribution of values that cell types show on different substrates. To ascertain potential underlying patterns in the distributional response of cell lines across varying ECM conditions, we used unsupervised machine learning to identify phenotypic classes (mechanotypes) for each of the physical features. These mechanotypes are characterized by the distribution of values the cells can attain given the cell type (defined by cell line in our case) and substrate conditions. Broader or narrower distribution of a particular mechanotype for a physical feature could be further viewed as the phenotypic plasticity of a cell type in terms of that particular physical feature.

Each cell line showed capabilities of switching between mechanotypes with change in substrate conditions for at least one of the physical features. This is particularly important in the context of cancer which is accompanied by extensive remodeling of the ECM^8^ and thus, depending on the local microenvironment cancer cells may equip themselves with means to attain required physical plasticity for fitness advantage. For example, consider the case of colon cancer cell lines, SW480 and SW620, that were established from the same patient, whereby SW480 was established from the primary site and SW620 was derived from a metastatic site^75^. While both show the same area mechanotype on most of the substrates, SW480 has a higher area mechanotype than SW620 on 30 kPa FN and Coll. They also show differing mechanotypes in terms of motility on 500 Pa Coll and 30 kPa Coll, and HA FN with SW620 having higher motility. SW620 also has a relatively softer cell mechanotype than SW480 across all the substrate conditions. Along similar lines, Tsujita et al. had reported that low-invasive breast cancer cells (MCF7) had higher plasma membrane tension as compared to highly metastatic breast cancer cells (MDA-MB-231)^76^. Consistent with this observation, we can see that MCF7 has a relatively stiffer mechanotype as compared to the MDA-MB-231 across substrates except on HA FN. Additionally, a clear distinction between breast normal and cancer cell lines can be seen in terms of cell stiffness, with normal cell lines having high cell stiffness mechanotype and cancer cell lines showing low to intermediate cell stiffness mechanotypes (Fig. 5D). Further, among the prostate metastatic cell lines a shift from low to high area mechanotype can be observed for low to highly metastatic^21,22^ (LNCaP to DU145 to PC-3) cell type for all except HA FN substrate. This analysis thus shows how the phenotypic plasticity in terms of physical characteristics of a cell type is inextricably linked to the ECM conditions. This is particularly important in the context of tumor progression, during which cancer cells experience substantial changes in the ECM mechanics and chemistry of their primary tissue and also encounter different ECM conditions in the metastatic tissue sites. Unraveling how cancer cells are able to exploit the ECM-based cues to alter their physical plasticity might provide therapeutic targets that could, for example, aid in abrogating cellular mechanical softness which helps cancer cells evade killing by cytotoxic T cells and thus, potentiate immunotherapy^77^.

In summary, this work underscores the fact that how cells process extracellular mechanical signals can be highly distinct between cell types or even for the same cell type on chemically distinct substrates leading to a wide range of physical responses. Future work will focus on relating the cellular microstates (characterized by key genes/proteins) underlying the mechanical and morphological responses to ECM-based cues. An important limitation of our study to note is that the pan-cancer mechanobiology dataset used here has been derived from subconfluent cells on a flat substrate. Differences from the observed behavior may arise due to other key ECM properties such as viscoelasticity^78,79^ and three dimensionality^80^. For example, glioblastoma cells in 3D agarose gels with collagen I reduce their migration speeds with increasing stiffness of the gel^81^, but the same cells on polyacrylamide gel with collagen I migrated faster with increasing stiffness of the gel^82^. Nevertheless, considering the lack of generalizability in cellular response to mechanical stimuli, future studies focusing on delineating how cells sense mechanical stimuli and how that is translated to phenotypic change should take into consideration the possibility of differing intrinsic characteristics (such as adhesion protein expression and type, molecular control of cytoskeletal assembly^83,84^, and vesicle trafficking for delivery and retrieval of transmembrane protein complexes^85^) leading to varied mechanosensitive responses, along with the augmentative role that chemical components of ECM such as HA can play in cell’s response.

## Methods

### Cell Growth

All cell lines were from ATCC (Manassas, VA) and grown according to ATCC standard operating procedures (SOPs), which has been deposited along with the data on figshare. Media and reagents used for cell culture were either provided by ATCC or purchased from an outside supplier as per ATCC SOP instructions. Thawed cells were taken to be P0. Cells were passaged until P3 and plated on polyacrylamide or HA gels at single cell density (25,000 cells/gel). Experiments were performed 24 hours after cell plating.

### Polyacrylamide Gel Fabrication

Polyacrylamide gels were created by using a combination of acrylamide and bisacrylamide from Bio-Rad Laboratories (Hercules, CA), along with water and NHS (N-hydroxysuccinimide ester) from Sigma (St. Louis, MO) dissolved in toluene (Fisher Scientific, Waltham, MA). Briefly, NHS was dissolved in toluene to create a saturated solution. The solution was then spun for 5 minutes on a tabletop centrifuge to remove any undissolved NHS. The NHS-toluene solution was then added to the water/polyacrylamide/bisacrylamide mixture and vortexed for ten seconds. To make the 30kPa polyacrylamide gels, a final concentration of 13.5% acrylamide was used, while 500 Pa polyacrylamide gels had a final concentration of 3.5%. Both gels had a bisacrylamide concentration of 0.33%. Acrylamide and bisacrylamide were from BioRad Laboratories (Hercules, CA). After vortexing the solution, we centrifuged at 1000 rpm in a tabletop centrifuge to separate the toluene from the polyacrylamide solution. We then removed the polyacrylamide solution from under the toluene layer, being careful not to disturb the toluene. We then aliquoted out the polyacrylamide. To begin polymerization we then added to a final concentration of 0.3% and 0.06% of Temed and APS (ammonium persulfate solution) respectively. Our solution was immediately pipetted onto functionalized coverslips and a siliconized coverslip was added on top. Gels were allowed to polymerize for 20 minutes. Top coverslips were removed and gels were rinsed twice for 5 minutes each in PBS. To crosslink protein to the surface of the gel, the gels were incubated in a 0.1mg/ml solution of collagen (BD Biosciences, San Jose, CA) or fibronectin (EMD Millipore, Billerica, MA) in HEPES pH 8.0 either overnight at 4 C or at RT for 4 hours. Gels were then rinsed and kept in PBS for storage.

### Hyaluronic Acid Gel Fabrication

HA gels were created by polymerizing thiol-modified hyaluronic acid with ExtraLink (PEGDA) (Ascendance Biotechnology, Alameda, CA) according to the manufacturer’s instructions. A 1ml vial of lyophilized HA was reconstituted with 875ul of degassed, deionized water for 30 minutes at 37C. After 30 minutes, 125ul of a 1mg/ml solution of either collagen or fibronectin was added to the vial. Gels were polymerized at a ratio of 1 part ExtraLink to 4 parts HA/protein solution. The desired volume of this mixture was pipetted on a glutaraldehyde-functionalized glass cover slip and then a siliconized coverslip was placed on top. Gels were allowed to polymerize for twenty minutes before removing the top coverslip and storing in PBS.

### Cell Seeding

To seed cells, gels in PBS were sterilized for 30 minutes under direct UV light. PBS was aspirated, warmed sterile media was added to the gels, and cells were added to single cell density (25,000 cells/18mm gel). Cells were allowed to adhere to the gels for 24 hours before experiments. After 24 hours, cells were imaged for morphology, single vs touching analysis, and motility. These analyses all used images taken on a Leica DMIRE2 microscope at 10x magnification. Also at 24 hours, cells were measured by AFM for stiffness.

### Cell morphology using light microscopy

Three metrics of cell morphology were determined using bright field imaging of live cells at 24 hours on each of the 7 substrates. Because of the need to image live cells without genetic or chemical perturbation, fluorescence microscopy cannot be used, and the contrast between cell and substrate, especially for soft gels, is too low to allow automated edge tracing algorithms to work. Therefore each cell was traced manually from images taken using a 10x lens and digital magnification to provide a pixel to micron ratio of 1.5. This resolution was optimal for allowing sufficient accuracy of tracing the cell contour while enabling simultaneous imaging of 5 to 10 single cells per field, based on our previous measurements using this method. Cells from two gel replicates per condition were traced, with up to 50 cells per replicate. The public domain NIH-developed software imageJ was used for tracing and to calculate three metrics: adherent area, aspect ratio, and circularity.

### Cell stiffness by Atomic Force Microscopy (AFM)

AFM measurements were conducted at room temperature using a Bioscope DAFMLN-AM head (Bruker, Santa Barbara, CA) mounted on an Axiovert 100 microscope (Zeiss, Thornwood, NY). The spring constants of triangular silicon nitride cantilevers with a 1µm diameter silica bead attached (Novascan, Ames, IA) were determined by resonance measurements (0.04 - 0.1 N/m). Indentations were made on three distinct areas of the cell avoiding the nucleus and the cell periphery. To quantify cellular stiffness (Young’s modulus), the first 500 nm of indentation into the cell was fit to the Hertz model for a sphere making contact with a homogenous, elastic half space

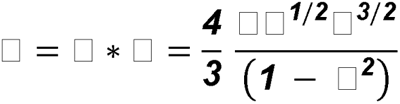

where *f* is the force applied to the cell, *k* is the spring constant of the cantilever, *d* is the deflection of the cantilever, *E* is the Young’s modulus, *R* is the radius of the bead, δ is the indentation into the cell and ν is the Poisson’s ratio of the cell (assumed to be 0.5). 10-15 cells were analyzed at 3 different points for each of the 7 substrate conditions per cell line.

### Motility

Cells were tracked, using the ImageJ cell tracker plugin, in time lapse movies made for periods between 1.75 and 3 hours taken after 24 hrs of incubation of the substrates. The centroid of each cell was tracked by localization of its nucleus at five-minute intervals. Speed was determined by dividing the total distance a cell moved by the time over which the cell was tracked.

### Unsupervised Clustering

The unsupervised clustering workflow was applied separately to each physical feature (area, aspect ratio, circularity, cell stiffness, and motility). Wasserstein-1 distance^25,26^ was used as the pairwise distance metric between cell line-substrate combinations and was computed using the R package maotai. The Wasserstein-1 distance measures the discrepancy between two distributions in terms of the area between their respective cumulative density functions (CDFs). Unlike Kullback-Lieibler (KL) divergence, a widely used measure for comparing probability distributions, Wasserstein-1 distance is symmetric and satisfies the triangular inequality, two fundamental properties for a distance to be considered as a metric. As compared to Jansen-Shannon divergence (symmetrized version of KL-divergence), Wasserstein-1 distance can be computed without estimating the probability density functions using kernel density estimates, thus avoiding the need to choose the bandwidth parameter of the smoothing kernel which could have significant effect on the clustering. Another distance metric often used to compare distribution is the Kolomogrov-Smirnov (KS) distance. The KS distance is defined as the largest absolute difference between the two empirical CDFs evaluated at any point and is bounded between 0 and 1. In contrast, Wasserstein-1 distance takes into consideration differences across the complete span of CDFs and is particularly good for distributions containing a significant amount of data in long tails. Thus, KS distance can level off due to its boundedness even if the distributions are considerably apart, whereas Wasserstein-1 distance will increase substantially which would aid in better separation of distinct distributions.

Consensus clustering method^23^ was used to determine the number and membership of possible classes for each physical feature. This unsupervised clustering method has been extensively used in cancer research to identify molecular subclasses based on omics data^86–88^. Clustering was performed using R package ConsensusClusterPlus^24^. In our analysis, the input to consensus clustering was the *n x n* matrix containing pairwise Wasserstein-1 distance between distributions, where *n* signifies the number of cell line-substrate pairs having at least 25 measurements. Note that a cell line-substrate pair corresponds to a particular cell line on a particular substrate. Consensus clustering involves repeated clustering of subsampled sets to determine the stability of a specified number of cluster count (*k*). For each of the physical features, we used 0.8 subsampling rate (i.e. 80% of *n* cell line-substrate pairs selected at random) and 1000 repetitions for cluster count *k* = 2, 3, …, 9. Partition around medoids (a form of k-medoids clustering method) was used to cluster the subsampled distance matrices. Then for each value of cluster count *k*, consensus matrix *M* is computed where element *M_ij_* is called consensus index for (*i^th^* cell line-substrate pair, *j^th^*cell line-substrate pair) defined as

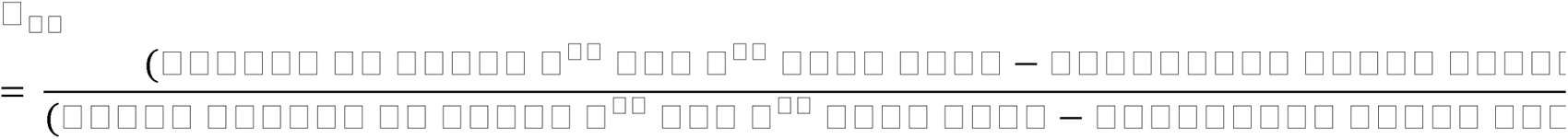

The optimal number of clusters, among the tested cluster counts *k* = 2, 3, …, 9, was determined using the proportion of ambiguous clustering (PAC)^27^ and Calinski-Harabasz Index (CHI)^28^. PAC corresponds to the fraction of *M_ij_* values between 0.1 and 0.9, which can be viewed as how many (*i^th^*cell line-substrate pair, *j^th^*cell line-substrate) pairs neither have high enough consensus (*M_ij_* > 0.9) to be considered in the same cluster nor low enough consensus (*M_ij_* < 0.1) to be considered in different clusters. CHI, on the other hand, compares the between-cluster separation with within-cluster dispersion. Thus, the optimal cluster count corresponds to low PAC and high CHI values. Hierarchical clustering can be used as a visualization tool to view a consensus matrix with high consensus index cell line-substrate pairs grouped together^23^. We used this to visualize the consensus matrices corresponding to the optimal cluster count (Supp. Figs. 7, 8).

Consensus indices can be used to item consensus values for each of the cell line-substrate pairs (items). Item consensus values correspond to the mean consensus of an item with all items in a particular cluster and thus, each cell line-substrate pair would have *k* item-consensus values corresponding to each cluster at a particular *k*. The cluster for which a cell line-substrate pair has the highest item-consensus value is considered its membership class. To further account for the inherent uncertainty in unsupervised clustering, after determining the optimal cluster count *k*, we identified the cell line-substrate pairs that do not show strong membership to its assigned class. That is, we considered the cell line-substrate pairs that have the highest item-consensus value of less than 0.8 as boundary cases. These boundary cases were instead viewed as being members of either of the two classes for which they have two largest item-consensus values (both less than 0.8) based on the available data. The same framework described here for Wasserstein-1 distance based clustering was used for KS distance based clustering. The heatmap visualizations for cluster membership of each cell line-substrate pair were created using ComplexHeatmap^89,90^.

### Statistical Analysis

Two-sided permutation test was used to assess statistical significance of the ratio of median values of a particular physical feature on two different substrate conditions being greater than 1, and of the difference between cancer cell lines and their normal counterpart(s) within the same tissue type in terms of the median value of the physical features. Benjamini-Hochberg procedure^91^ was used to adjust the p-values for multiple testing.

## Data Availability

All the measurement data for morphology (area, aspect ratio, and circularity), cell stiffness, and cell speed used in this study has been deposited at 10.6084/m9.figshare.27927585.

## Code Availability

All analyses reported in this study used the statistical software R (v.4.4.2). All R files are available publicly at https://github.com/kparihar13/pson_cell_line_mechanobiology_analysis.

## Acknowledgements

The data was obtained with support from the National Cancer Institute and Leidos Biomedical Research, Inc. under contract 15X008 with Frederick National Laboratory for Cancer Research. This study has also received funding from the National Institutes of Health under R35GM136259 and U01CA250044. Figures 1A and 5A were created with BioRender.com.

## Author Contributions

PAJ designed the experimental mechanobiology research. KC managed the experimental research project. KC, LC, MEM, MGM, ASvO, AH, EEC, PAG, MD, and TL performed cell culture and imaging experiments. FJB performed atomic force microscopy measurements. DVI performed experiments and assisted in data interpretation. KP, JN, PAJ, and RR designed the data analysis and machine learning research. KP performed statistical analysis and machine learning. KP, PAJ, and RR wrote the article.

## Competing Interests

The authors declare no competing interests.

## Notes

https://github.com/kparihar13/pson_cell_line_mechanobiology_analysis

### Competing Interest Statement

The authors have declared no competing interest.

