## Supplementary material for "Tissue-dependent mechanosensing by cells derived from human tumors": SI Tables and Figures

Table 1: Number of single cell measurements of spread area, aspect ratio, and circularity for each cell line-substrate combination.

| | 30kPa Coll | 30kPa FN | 500Pa Coll | 500Pa FN | Glass | HA Coll | HA FN |
|---|---|---|---|---|---|---|---|
| SK-MEL-2 | 100 | 100 | 101 | 100 | 100 | 101 | 100 |
| A375 | 100 | 100 | 101 | 100 | 100 | 100 | 100 |
| WM266-4 | 100 | 100 | 100 | 100 | 100 | 100 | 100 |
| MeWo | 50 | 50 | 100 | 96 | 100 | 100 | 80 |
| RWPE-1(N) | 102 | 103 | 100 | 103 | 100 | 101 | 101 |
| 22Rv1 | 99 | 100 | 99 | 81 | 103 | 103 | 98 |
| LnCaP | 41 | 49 | 34 | 59 | 90 | 32 | 46 |
| DU145 | 77 | 99 | 67 | 101 | 81 | 102 | 103 |
| PC-3 | 101 | 100 | 100 | 90 | 100 | 100 | 40 |
| hTERT-HPNE(N) | 94 | 100 | 106 | 97 | 100 | 77 | 76 |
| Panc-1 | 100 | 102 | 72 | 21 | 101 | 103 | 105 |
| Capan-1 | 105 | 100 | 102 | 100 | 101 | 101 | 100 |
| SKOV-3 | 108 | 101 | 82 | 104 | 102 | 100 | 92 |
| Caov-3 | 85 | 34 | 37 | 16 | 81 | 59 | 43 |
| OVCAR-3 | 100 | 86 | 94 | 89 | 90 | 100 | 97 |
| NL20(N) | 168 | 257 | 247 | 284 | 248 | 126 | 150 |
| NCI-H2126 | 100 | 100 | 100 | 100 | 100 | 100 | 100 |
| NCI-H2087 | 100 | 100 | 100 | 100 | 100 | 100 | 100 |
| HCT116 | 100 | 100 | 107 | 101 | 101 | 100 | 100 |
| HT29 | 82 | 55 | 93 | 40 | 165 | 95 | 49 |
| SW480 | 100 | 100 | 100 | 100 | 100 | 100 | 100 |
| SW620 | 101 | 100 | 100 | 100 | 101 | 100 | 100 |
| hTERT-HME1(N) | 100 | 100 | 101 | 100 | 100 | 74 | 51 |
| MCF10A(N) | 100 | 100 | 100 | 100 | 91 | 88 | 62 |
| T-47D | 100 | 100 | 100 | 86 | 100 | 84 | 97 |
| MCF7 | 100 | 68 | 100 | 82 | 100 | 45 | 29 |
| MDA-MB-231 | 121 | 218 | 145 | 164 | 193 | 77 | 160 |
| HCC1937 | 100 | 100 | 100 | 100 | 100 | 100 | 100 |
| U-87 | 101 | 101 | 102 | 100 | 101 | 101 | 100 |
| T98G | 100 | 101 | 100 | 101 | 101 | 100 | 101 |

Table 2: Number of measurements of cell stiffness for each cell line-substrate combination.

| | 30kPa Coll | 30kPa FN | 500Pa Coll | 500Pa FN | Glass | HA Coll | HA FN |
|---|---|---|---|---|---|---|---|
| SK-MEL-2 | 45 | 44 | 43 | 47 | 45 | 50 | 43 |
| A375 | 45 | 30 | 39 | 44 | 45 | 42 | 48 |
| WM266-4 | 63 | 44 | 42 | 41 | 44 | 41 | 48 |
| MeWo | 29 | 31 | 30 | 33 | 30 | 33 | 30 |
| RWPE-1(N) | 30 | 36 | 43 | 48 | 67 | 45 | 30 |
| 22Rv1 | 30 | 30 | 45 | 30 | 45 | 42 | 30 |
| LnCaP | 29 | 15 | 27 | 12 | 48 | NA | 38 |
| DU145 | 47 | 44 | 51 | 51 | 49 | 41 | 50 |
| PC-3 | 45 | 39 | 35 | 29 | 45 | 30 | 28 |
| hTERT-HPNE(N) | 45 | 49 | 50 | 42 | 45 | 45 | 45 |
| Panc-1 | 36 | 43 | 30 | 39 | 45 | 43 | 44 |
| Capan-1 | 44 | 31 | 29 | 27 | 33 | 32 | 29 |
| SKOV-3 | 44 | 39 | 29 | 27 | 28 | 30 | 26 |
| Caov-3 | 44 | 45 | 44 | 45 | 45 | 44 | 28 |
| OVCAR-3 | 45 | 44 | 45 | 44 | 46 | 47 | 45 |
| NL20(N) | 30 | 30 | 30 | 33 | 33 | 30 | 32 |
| NCI-H2126 | 30 | 45 | 29 | 30 | 30 | 30 | 30 |
| NCI-H2087 | 30 | 29 | 32 | 30 | 30 | 51 | 30 |
| HCT116 | 65 | 45 | 49 | 42 | 67 | 66 | 54 |
| HT29 | 45 | 39 | 45 | 32 | 45 | 45 | 48 |
| SW480 | 44 | 53 | 45 | 59 | 63 | 45 | 45 |
| SW620 | 31 | 29 | 45 | 32 | 31 | 30 | 48 |
| hTERT-HME1(N) | 45 | 45 | 30 | 47 | 47 | 27 | 45 |
| MCF10A(N) | 45 | 44 | 44 | 45 | 60 | 45 | 45 |
| T-47D | 48 | 44 | 45 | 45 | 48 | 39 | 42 |
| MCF7 | 52 | 42 | 47 | 42 | 29 | 45 | 45 |
| MDA-MB-231 | 45 | 45 | 30 | 30 | 32 | 30 | 45 |
| HCC1937 | 42 | 30 | 45 | 44 | 39 | 42 | 42 |
| U-87 | 51 | 35 | 59 | 54 | 72 | 54 | 45 |
| T98G | 63 | 64 | 59 | 61 | 61 | 57 | 56 |

Table 3: Number of single cell measurements of cell speed for each cell line-substrate combination.

| | 30kPa Coll | 30kPa FN | 500Pa Coll | 500Pa FN | Glass | HA Coll | HA FN |
|---|---|---|---|---|---|---|---|
| SK-MEL-2 | 71 | 72 | 61 | 141 | 129 | 70 | 68 |
| A375 | 50 | 50 | 50 | 50 | 51 | 52 | 51 |
| WM266-4 | 41 | 46 | 54 | 51 | 50 | 51 | 51 |
| MeWo | 176 | 58 | 255 | 53 | 159 | 30 | 72 |
| RWPE-1(N) | 119 | 109 | 114 | 102 | 66 | 108 | 105 |
| 22Rv1 | 36 | 42 | 50 | 45 | 49 | 50 | 56 |
| LnCaP | 48 | 50 | 29 | 50 | 50 | 48 | 53 |
| DU145 | 41 | 11 | 27 | 43 | 32 | 50 | 40 |
| PC-3 | 13 | 21 | 13 | 99 | 101 | 24 | 14 |
| hTERT-HPNE(N) | 55 | 52 | 51 | 51 | 52 | 47 | 50 |
| Panc-1 | 45 | 44 | 33 | 8 | 57 | 62 | 55 |
| Capan-1 | 52 | 46 | 79 | 17 | 56 | 53 | 30 |
| SKOV-3 | 33 | 26 | 39 | 49 | 51 | 62 | 44 |
| Caov-3 | 22 | 17 | 25 | 4 | 24 | 30 | 27 |
| OVCAR-3 | 29 | 52 | 49 | 53 | 58 | 51 | 50 |
| NL20(N) | 61 | 70 | 54 | 67 | 26 | 45 | 44 |
| NCI-H2126 | 51 | 56 | 72 | 75 | 85 | 56 | 103 |
| NCI-H2087 | 87 | 27 | 103 | 114 | 104 | 93 | 69 |
| HCT116 | 47 | 36 | 44 | 9 | 70 | 26 | 23 |
| HT29 | 50 | 36 | 57 | 15 | 50 | 50 | 50 |
| SW480 | 56 | 50 | 55 | 83 | NA | 40 | 65 |
| SW620 | 25 | 25 | 51 | 51 | 25 | 51 | 36 |
| hTERT-HME1(N) | 51 | 29 | 50 | 54 | 52 | 20 | 32 |
| MCF10A(N) | 50 | 50 | 50 | 60 | 50 | 50 | 52 |
| T-47D | 30 | 56 | 31 | 47 | 15 | 81 | 27 |
| MCF7 | 22 | 15 | 37 | 3 | 50 | 50 | 36 |
| MDA-MB-231 | 27 | 25 | 30 | 29 | 34 | 12 | 26 |
| HCC1937 | 51 | 49 | 54 | 50 | 50 | 53 | 48 |
| U-87 | 59 | 54 | 52 | 102 | 70 | 58 | 47 |
| T98G | 74 | 82 | 123 | 94 | 62 | 75 | 86 |

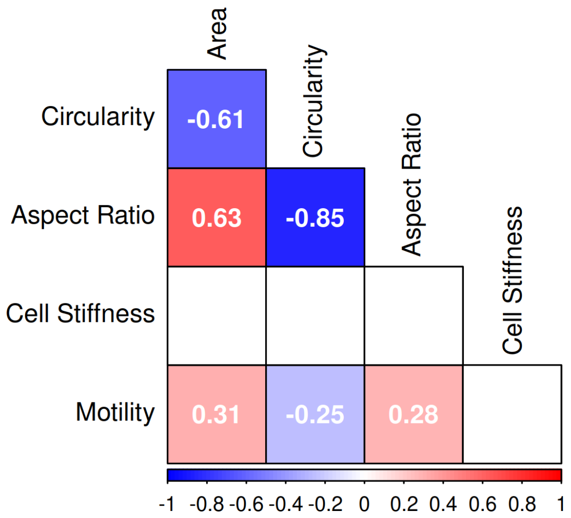


Supplementary Figure 1: Spearman correlation matrix using the median values of the phenotypic features for the cell line-substrate pairs. Only statistically significant correlations are shown. Note that in this analysis only those cell line-substrate pairs which have at least 25 data points for each of the physical feature are considered.

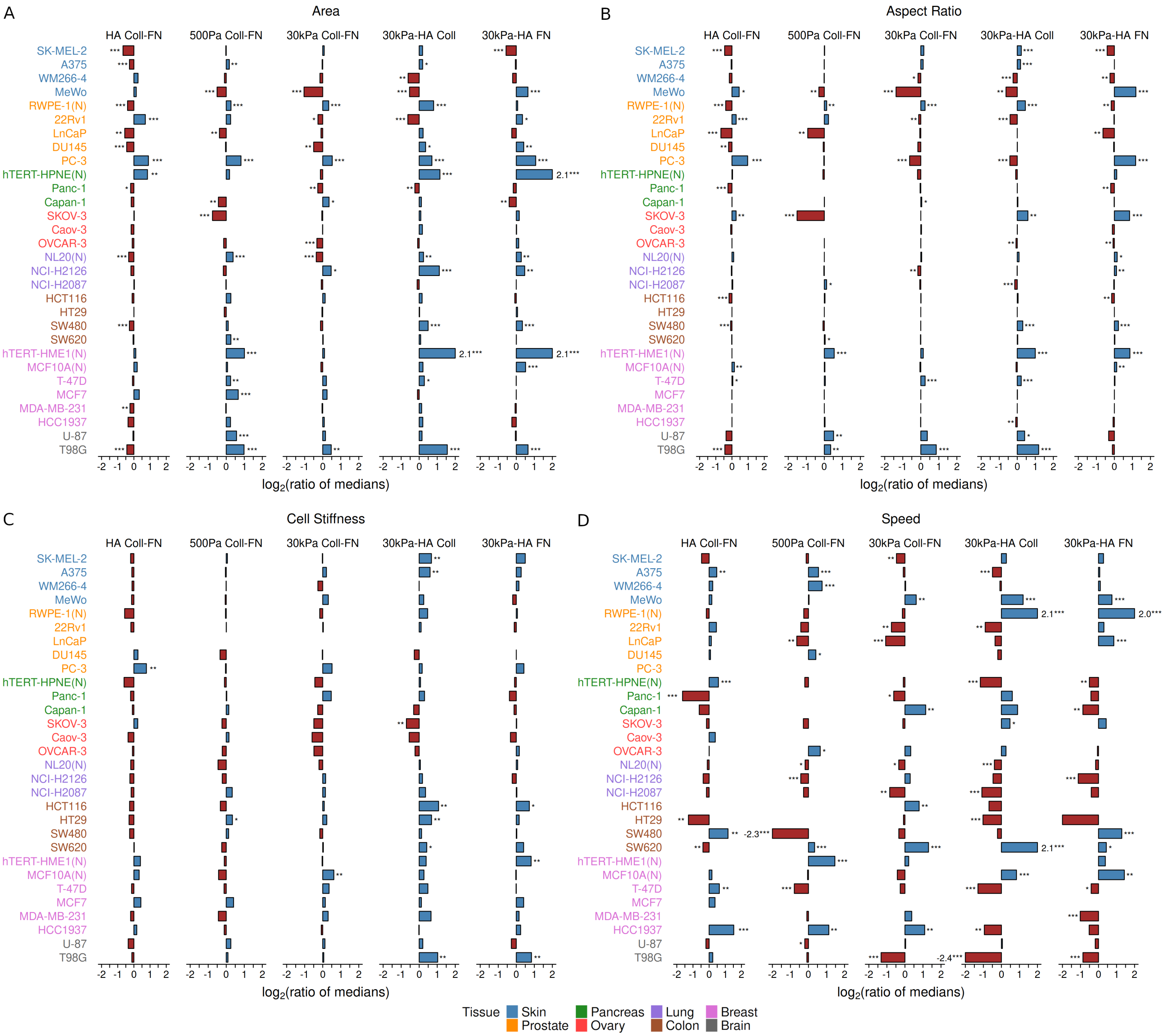


Supplementary Figure 2: For each cell line, the ratio of the median values for (A) area, (B) shape (aspect ratio), (C) cell stiffness, and (D) speed as a measure of the phenotypic sensitivity to substrate change. (HA Coll-FN: HA Coll/HA FN, 500Pa Coll-FN: 500Pa FN/500Pa Coll, 30kPa Coll-FN: 30kPa Coll/30kPa FN, 30kPa-HA Coll: 30kPa Coll/HA Coll, 30kPa-HA FN: 30kPa FN/HA FN). (N) refers to non-malignant (normal) cell lines. ***p-value $< 0.01$; **p-value $< 0.05$; *p-value $< 0.1$. For each cell line on a particular substrate, $n > 25$ cells. See supplementary tables 1-3 for the exact value of $n$ for the cell lines.

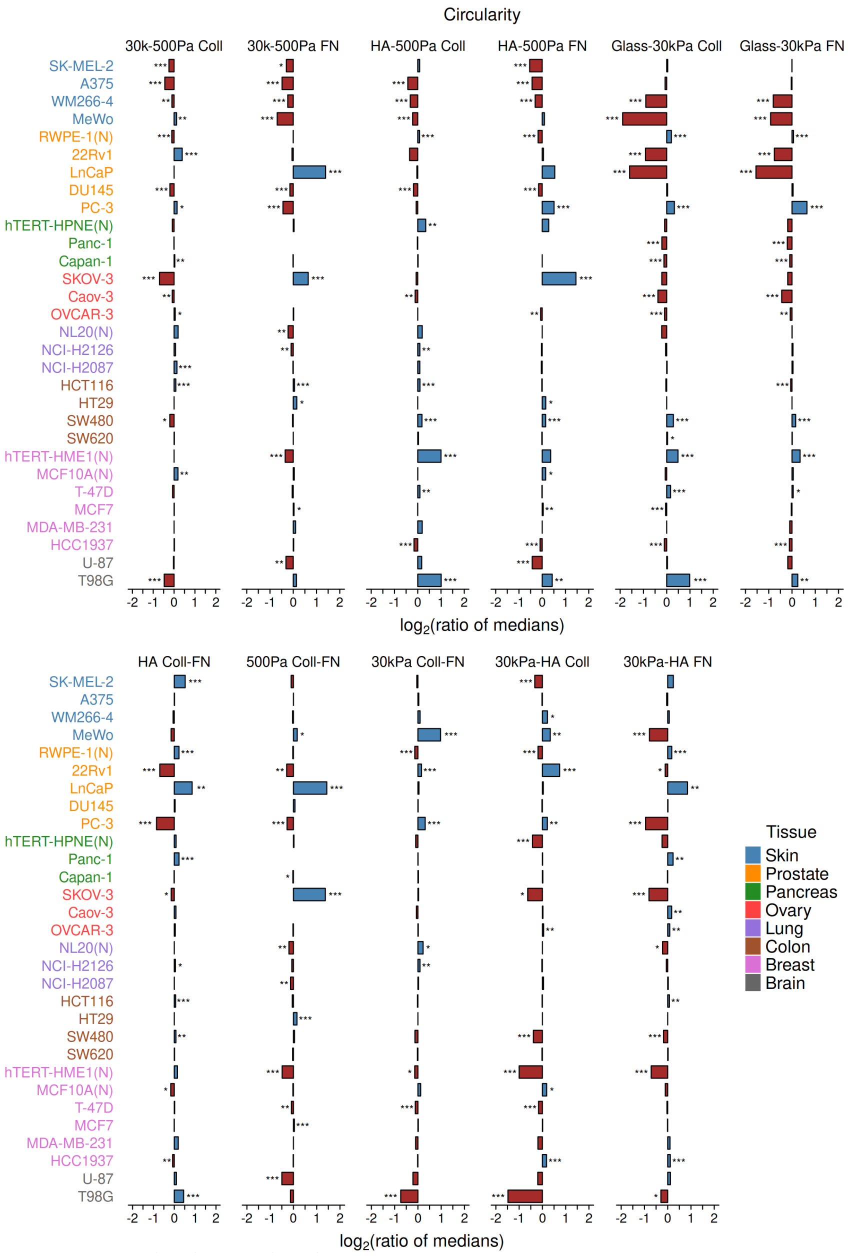


Supplementary Figure 3: For each cell line, the ratio of the median values for circularity as a measure of the phenotypic sensitivity to substrate change. (30k-500Pa Coll: 30kPa Coll/500Pa Coll, 30k-500Pa FN: 30kPa FN/500Pa FN, HA-500Pa Coll: HA Coll/500Pa Coll, HA-500Pa FN: HA FN/500Pa FN, Glass-30kPa Coll: Glass/30kPa Coll, Glass-30kPa FN: Glass/30kPa FN, HA Coll-FN: HA Coll/HA FN, 500Pa Coll-FN: 500Pa FN/500Pa Coll, 30kPa Coll-FN: 30kPa Coll/30kPa FN, 30kPa-HA Coll: 30kPa Coll/HA Coll, 30kPa-HA FN: 30kPa FN/HA FN). (N) refers to non-malignant (normal) cell lines. ***p-value $< 0.01$; **p-value $< 0.05$; *p-value $< 0.1$. For each cell line on a particular substrate, $n > 25$ cells. See supplementary tables 1-3 for the exact value of $n$ for the cell lines.

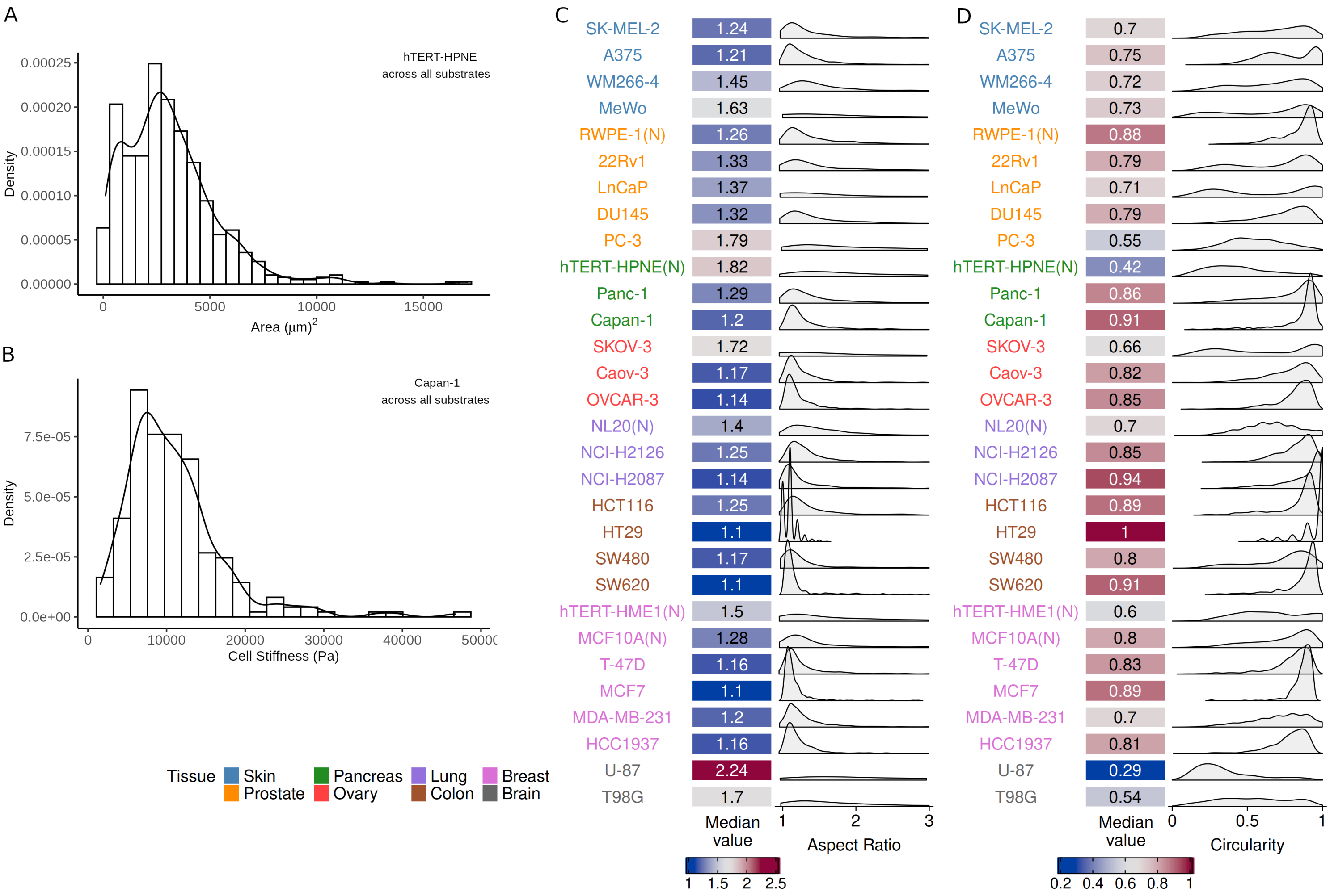


Supplementary Figure 4: KDEs for outlier-like distributions of (A) hTERT-HPNE cellular area across all substrates, and (B) Capan-1 cell stiffness across all substrates. Median values of cell (A) aspect ratio, and (B) circularity across all the 7 different substrates, along with the kernel density estimates (KDE) for summarizing the distribution of feature values for each cell line. (N) refers to non-malignant (normal) cell lines. Measurements from $n > 200$ cells for each of the cell lines. See supplementary tables 1-3 for the exact value of $n$ for the cell lines.

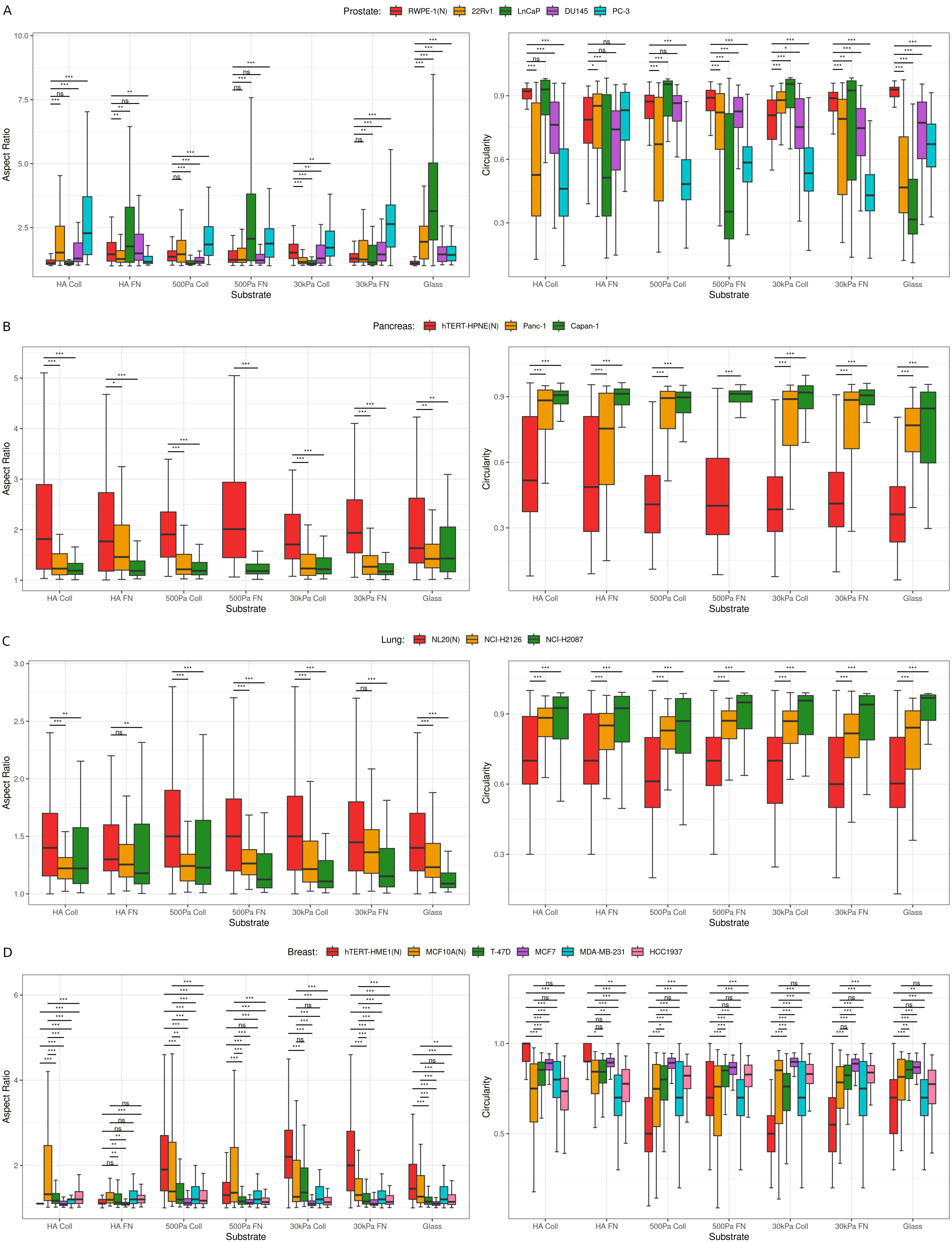
A
Prostate: RWPE-1(N) 22Rv1 LnCaP DU145 PC-3
Aspect Ratio
Circularity
Substrate
HA Coll HA FN 500Pa Coll 500Pa FN 30kPa Coll 30kPa FN Glass
B
Pancreas: hTERT-HPNE(N) Panc-1 Capan-1
Aspect Ratio
Circularity
Substrate
HA Coll HA FN 500Pa Coll 500Pa FN 30kPa Coll 30kPa FN Glass
C
Lung: NL20(N) NCI-H2126 NCI-H2087
Aspect Ratio
Circularity
Substrate
HA Coll HA FN 500Pa Coll 500Pa FN 30kPa Coll 30kPa FN Glass
D
Breast: hTERT-HME1(N) MCF10A(N) T-47D MCF7 MDA-MB-231 HCC1937
Aspect Ratio
Circularity
Substrate
HA Coll HA FN 500Pa Coll 500Pa FN 30kPa Coll 30kPa FN Glass

Supplementary Figure 5: Comparing tissue-specific normal and cancer cell behavior in terms of aspect ratio, and cicularity for (A) prostate, (B) panacrease, (C) lung, and (D) breast cell lines soft (500 Pa) and stiff (30 kPa) PAAm Coll and FN substrates, soft (500 Pa) HA substrates coated with Coll and FN, and glass. (N) refers to non-malignant (normal) cell lines. ***p-value $< 0.01$; **p-value $< 0.05$; *p-value $< 0.1$. For each cell line on a particular substrate, $n > 25$ cells. See supplementary tables 1-3 for the exact value of $n$ for the cell lines.

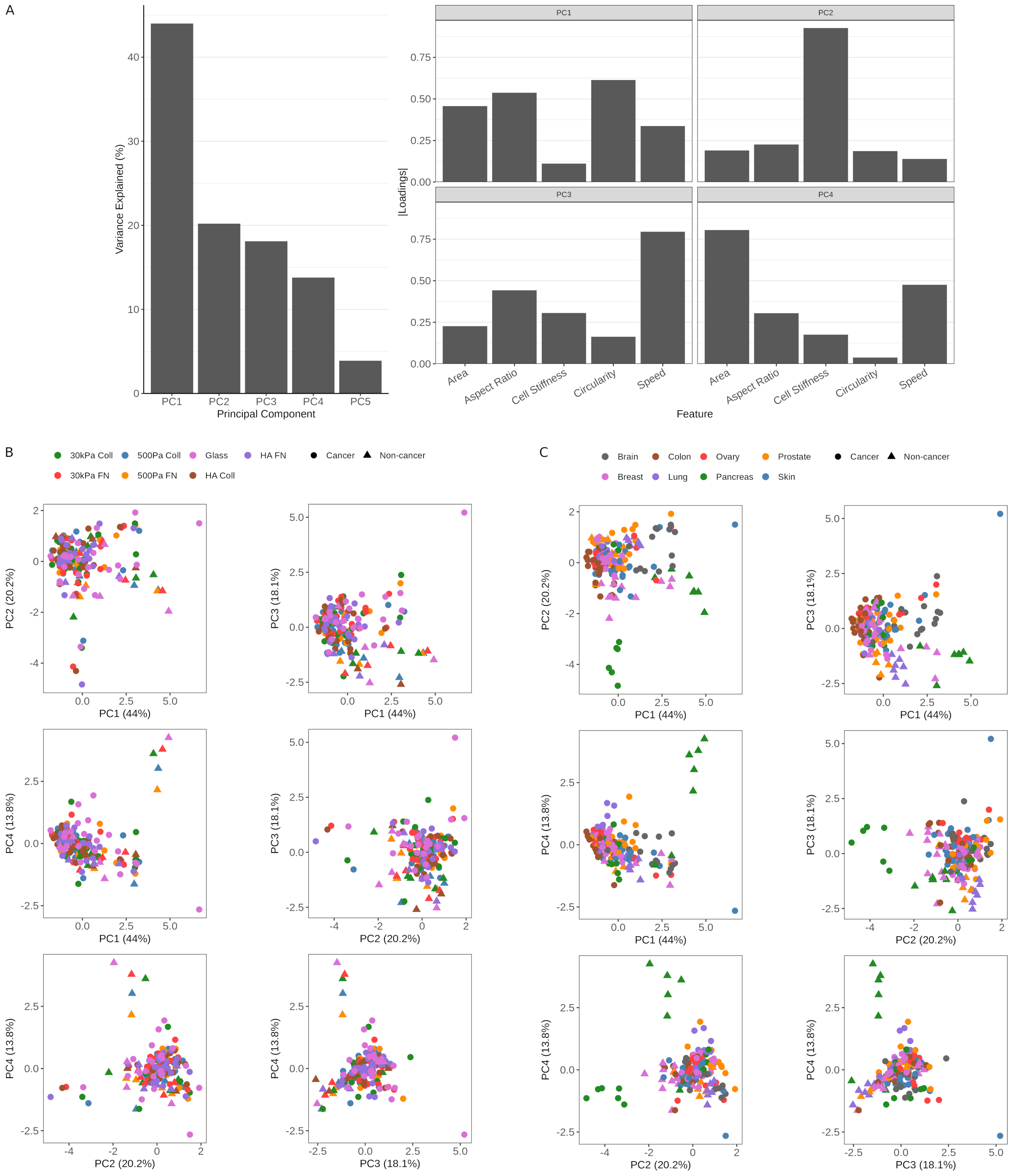
A
Variance Explained (%)
Principal Component
PC1
PC2
PC3
PC4
PC5
|Loadings|
Feature
Area
Aspect Ratio
Cell Stiffness
Circularity
Speed
B
30kPa Coll
500Pa Coll
Glass
HA FN
30kPa FN
500Pa FN
HA Coll
Cancer
Non-cancer
PC1 (44%)
PC2 (20.2%)
PC3 (18.1%)
PC4 (13.8%)
C
Brain
Colon
Ovary
Prostate
Breast
Lung
Pancreas
Skin

Supplementary Figure 6: Principal component analysis (PCA) performed using the median values of cell area, aspect ratio, circularity, stiffness, and speedfrom all the cell line-substrate pairs. Note that in this analysis only those cell line-substrate pairs which have at least 25 data points for each of the physical feature are considered. (A) Vaiance explained by each of the PCs, and the loadings of the physical features on the first four PCs. Pairwise scatter plots for the first four PCs, with the points colored by (B) the substrate type and (C) tissue type.

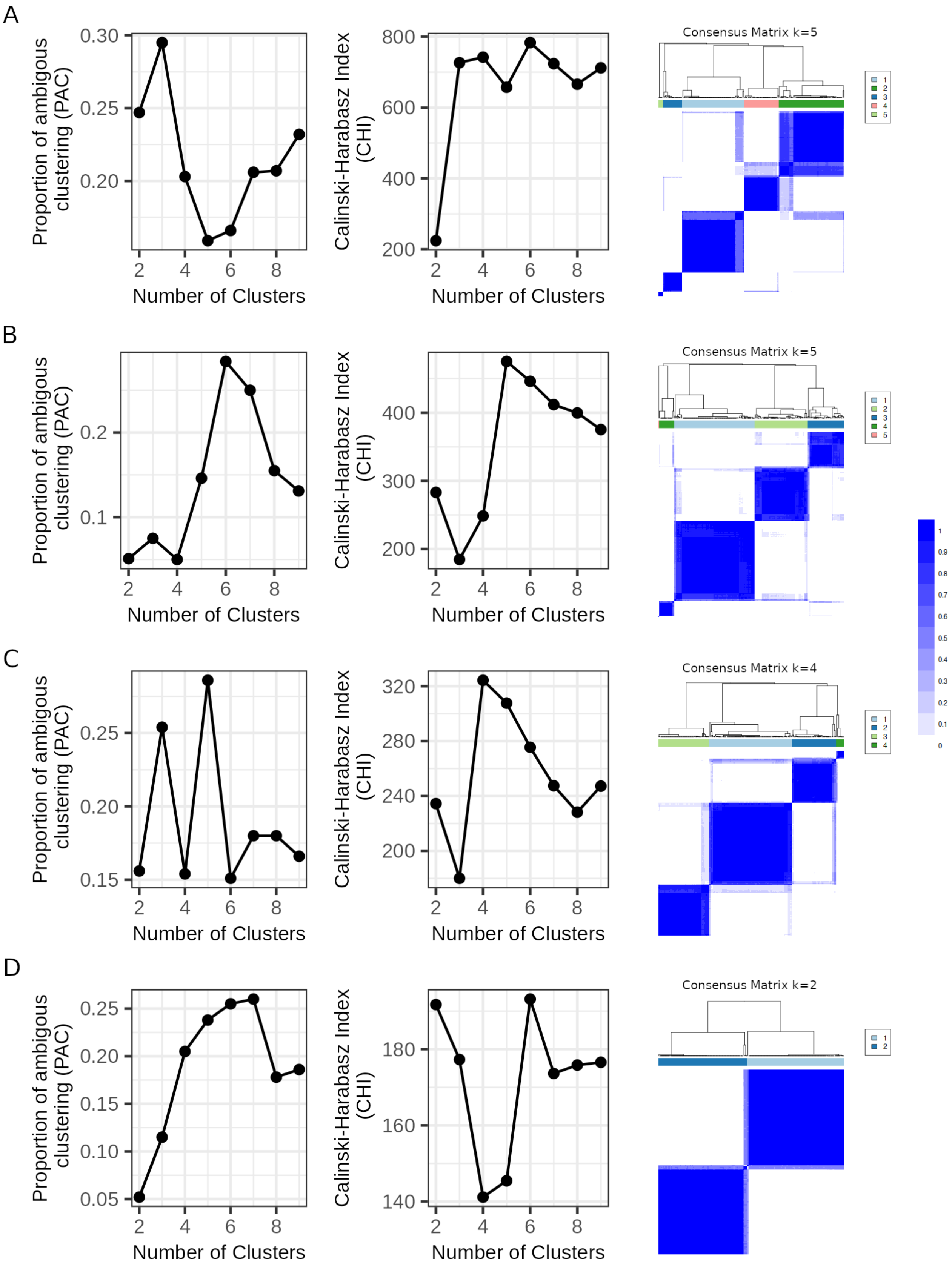


Supplementary Figure 7: Clustering statistics (PAC and CHI) used for identifying the optimal number of clusters (mechanotypes) based on Wasserstein-1 distance and the corresponding consensus matrix for optimal cluster count $k$ of cell (A) area, (B) aspect ratio, (C) stiffness, and (D) speed.

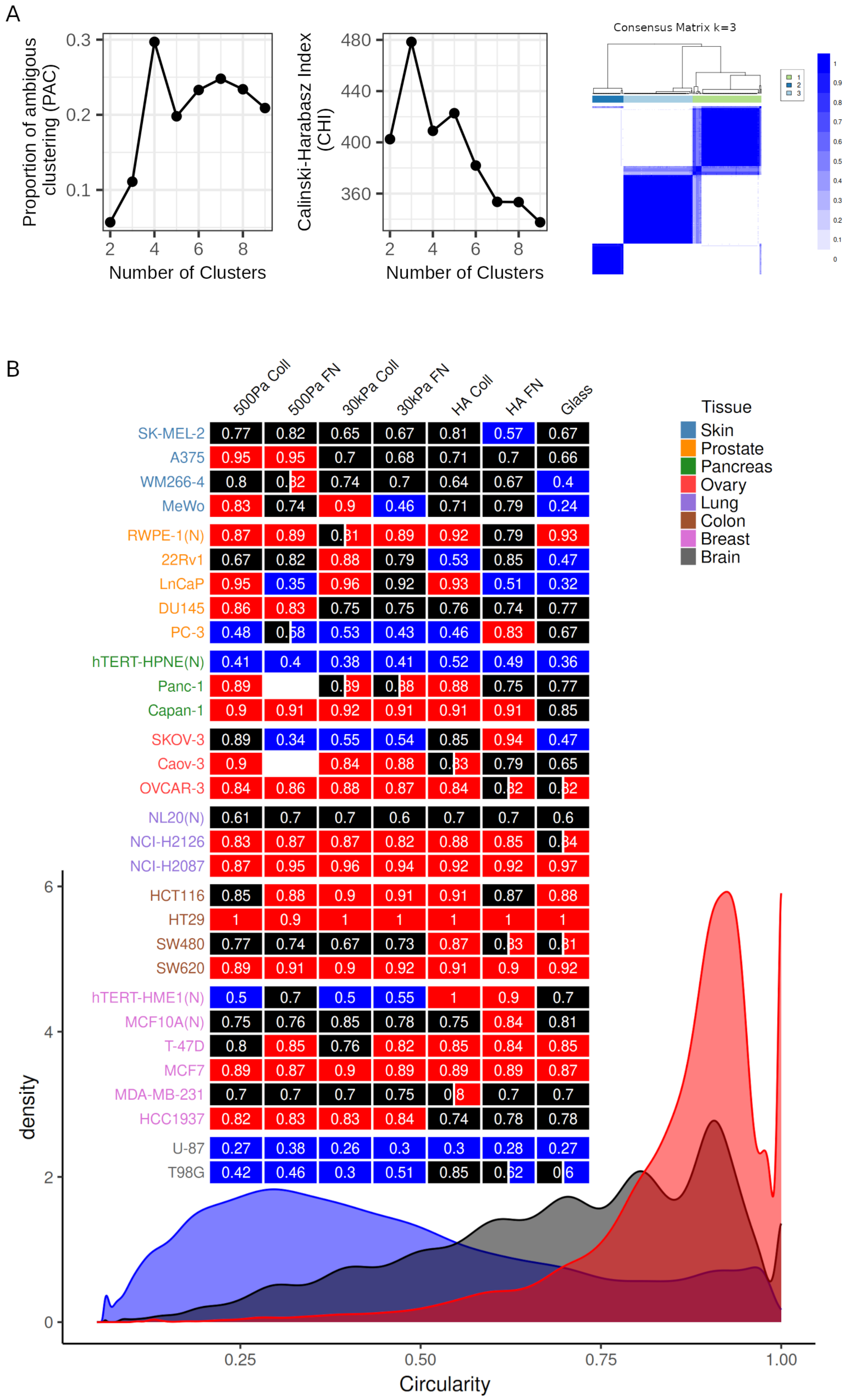
A
Proportion of ambiguous clustering (PAC)
0.3
0.2
0.1
2 4 6 8
Number of Clusters
Calinski-Harabasz Index (CHI)
480
440
400
360
2 4 6 8
Number of Clusters
Consensus Matrix k=3
1
2
3
B
500Pa Coll 500Pa FN 30kPa Coll 30kPa FN HA Coll HA FN Glass
SK-MEL-2 0.77 0.82 0.65 0.67 0.81 0.57 0.67
A375 0.95 0.95 0.7 0.68 0.71 0.7 0.66
WM266-4 0.8 0.32 0.74 0.7 0.64 0.67 0.4
MeWo 0.83 0.74 0.9 0.46 0.71 0.79 0.24
RWPE-1(N) 0.87 0.89 0.31 0.89 0.92 0.79 0.93
22Rv1 0.67 0.82 0.88 0.79 0.53 0.85 0.47
LnCaP 0.95 0.35 0.96 0.92 0.93 0.51 0.32
DU145 0.86 0.83 0.75 0.75 0.76 0.74 0.77
PC-3 0.48 0.58 0.53 0.43 0.46 0.83 0.67
hTERT-HPNE(N) 0.41 0.4 0.38 0.41 0.52 0.49 0.36
Panc-1 0.89 0.39 0.38 0.88 0.75 0.77
Capan-1 0.9 0.91 0.92 0.91 0.91 0.91 0.85
SKOV-3 0.89 0.34 0.55 0.54 0.85 0.94 0.47
Caov-3 0.9 0.84 0.88 0.33 0.79 0.65
OVCAR-3 0.84 0.86 0.88 0.87 0.84 0.32 0.32
NL20(N) 0.61 0.7 0.7 0.6 0.7 0.7 0.6
NCI-H2126 0.83 0.87 0.87 0.82 0.88 0.85 0.34
NCI-H2087 0.87 0.95 0.96 0.94 0.92 0.92 0.97
HCT116 0.85 0.88 0.9 0.91 0.91 0.87 0.88
HT29 1 0.9 1 1 1 1 1
SW480 0.77 0.74 0.67 0.73 0.87 0.33 0.31
SW620 0.89 0.91 0.9 0.92 0.91 0.9 0.92
hTERT-HME1(N) 0.5 0.7 0.5 0.55 1 0.9 0.7
MCF10A(N) 0.75 0.76 0.85 0.78 0.75 0.84 0.81
T-47D 0.8 0.85 0.76 0.82 0.85 0.84 0.85
MCF7 0.89 0.87 0.9 0.89 0.89 0.89 0.87
MDA-MB-231 0.7 0.7 0.7 0.75 0.8 0.7 0.7
HCC1937 0.82 0.83 0.83 0.84 0.74 0.78 0.78
U-87 0.27 0.38 0.26 0.3 0.3 0.28 0.27
T98G 0.42 0.46 0.3 0.51 0.85 0.62 0.6
Tissue
Skin
Prostate
Pancreas
Ovary
Lung
Colon
Breast
Brain
density
6
4
2
0
0.25 0.50 0.75 1.00
Circularity

Supplementary Figure 8: (A) Clustering statistics (PAC and CHI) used for identifying the optimal number of clusters (mechanotypes) based on Wasserstein-1 distance and the corresponding consensus matrix for optimal cluster count $k$ of circularity. (B) Mechanotypes for circularity, whereby the heatmap shows the phenotypic class for each cell line-substrate pair and the KDEs correspond to characteristic density function for each class. The numeric values shown in the heatmap correspond to median values of circularity for each cell line-substrate pair. (N) refers to non-malignant (normal) cell lines. Note that in this analysis only those cell line-substrate pairs which have at least 25 data points for the physical feature of interest are considered.

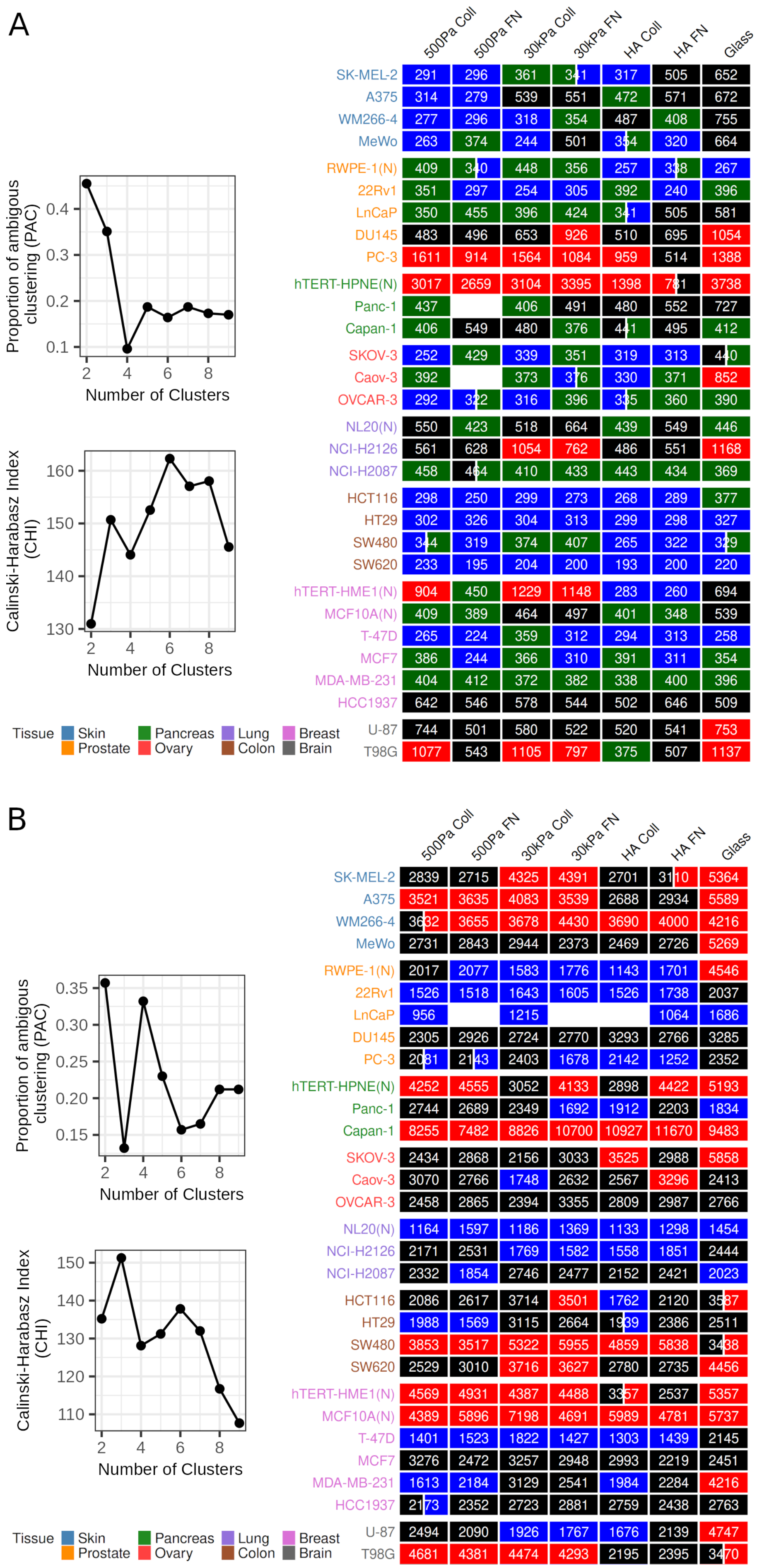
A
Proportion of ambigous clustering (PAC)
0.1 0.2 0.3 0.4
2 4 6 8
Number of Clusters
Calinski-Harabasz Index (CHI)
130 140 150 160
2 4 6 8
Number of Clusters
500Pa Coll 500Pa FN 30kPa Coll 30kPa FN HA Coll HA FN Glass
SK-MEL-2 291 296 361 341 317 505 652
A375 314 279 539 551 472 571 672
WM266-4 277 296 318 354 487 408 755
MeWo 263 374 244 501 354 320 664
RWPE-1(N) 409 340 448 356 257 338 267
22Rv1 351 297 254 305 392 240 396
LnCaP 350 455 396 424 341 505 581
DU145 483 496 653 926 510 695 1054
PC-3 1611 914 1564 1084 959 514 1388
hTERT-HPNE(N) 3017 2659 3104 3395 1398 781 3738
Panc-1 437 406 491 480 552 727
Capan-1 406 549 480 376 441 495 412
SKOV-3 252 429 339 351 319 313 440
Caov-3 392 373 376 330 371 852
OVCAR-3 292 322 316 396 335 360 390
NL20(N) 550 423 518 664 439 549 446
NCI-H2126 561 628 1054 762 486 551 1168
NCI-H2087 458 464 410 433 443 434 369
HCT116 298 250 299 273 268 289 377
HT29 302 326 304 313 299 298 327
SW480 344 319 374 407 265 322 329
SW620 233 195 204 200 193 200 220
hTERT-HME1(N) 904 450 1229 1148 283 260 694
MCF10A(N) 409 389 464 497 401 348 539
T-47D 265 224 359 312 294 313 258
MCF7 386 244 366 310 391 311 354
MDA-MB-231 404 412 372 382 338 400 396
HCC1937 642 546 578 544 502 646 509
U-87 744 501 580 522 520 541 753
T98G 1077 543 1105 797 375 507 1137
Tissue Skin Prostate Pancreas Ovary Lung Colon Breast Brain
B
Proportion of ambigous clustering (PAC)
0.15 0.20 0.25 0.30 0.35
2 4 6 8
Number of Clusters
Calinski-Harabasz Index (CHI)
110 120 130 140 150
2 4 6 8
Number of Clusters
500Pa Coll 500Pa FN 30kPa Coll 30kPa FN HA Coll HA FN Glass
SK-MEL-2 2839 2715 4325 4391 2701 3110 5364
A375 3521 3635 4083 3539 2688 2934 5589
WM266-4 3632 3655 3678 4430 3690 4000 4216
MeWo 2731 2843 2944 2373 2469 2726 5269
RWPE-1(N) 2017 2077 1583 1776 1143 1701 4546
22Rv1 1526 1518 1643 1605 1526 1738 2037
LnCaP 956 1215 1064 1686
DU145 2305 2926 2724 2770 3293 2766 3285
PC-3 2081 2143 2403 1678 2142 1252 2352
hTERT-HPNE(N) 4252 4555 3052 4133 2898 4422 5193
Panc-1 2744 2689 2349 1692 1912 2203 1834
Capan-1 8255 7482 8826 10700 10927 11670 9483
SKOV-3 2434 2868 2156 3033 3525 2988 5858
Caov-3 3070 2766 1748 2632 2567 3296 2413
OVCAR-3 2458 2865 2394 3355 2809 2987 2766
NL20(N) 1164 1597 1186 1369 1133 1298 1454
NCI-H2126 2171 2531 1769 1582 1558 1851 2444
NCI-H2087 2332 1854 2746 2477 2152 2421 2023
HCT116 2086 2617 3714 3501 1762 2120 3587
HT29 1988 1569 3115 2664 1939 2386 2511
SW480 3853 3517 5322 5955 4859 5838 3438
SW620 2529 3010 3716 3627 2780 2735 4456
hTERT-HME1(N) 4569 4931 4387 4488 3357 2537 5357
MCF10A(N) 4389 5896 7198 4691 5989 4781 5737
T-47D 1401 1523 1822 1427 1303 1439 2145
MCF7 3276 2472 3257 2948 2993 2219 2451
MDA-MB-231 1613 2184 3129 2541 1984 2284 4216
HCC1937 2173 2352 2723 2881 2759 2438 2763
U-87 2494 2090 1926 1767 1676 2139 4747
T98G 4681 4381 4474 4293 2195 2395 3470
Tissue Skin Prostate Pancreas Ovary Lung Colon Breast Brain

Supplementary Figure 9: Clustering statistics (PAC and CHI) used for identifying the optimal number of clusters based on Kolomogrov-Smirnov distance and the heatmap showing the corresponding classes for each of the cell line-substrate pairs for cell (A) area, and (B) stiffness. The numeric values shown in the heatmap correspond to median values of circularity for each cell line-substrate pair. (N) refers to non-malignant (normal) cell lines. Note that in this analysis only those cell line-substrate pairs which have at least 25 data points for the physical feature of interest are considered.
